# A sterol reductase responsible for the unusual 8(14)-unsaturation in bacterial sterol production and degradation

**DOI:** 10.64898/2026.02.26.708318

**Authors:** Alysha K. Lee, José-Luis Giner, Paula V. Welander

**Author notes:** Address correspondence to Paula V. Welander.

## Abstract

Sterols are a class of lipids that play a crucial role in human health through their essential physiological roles and as a point of interaction between commensal and pathogenic bacteria. The biosynthesis and modification of these lipids is a well-characterized process in many eukaryotes and increasingly in bacteria. However, the proteins responsible for formation of the unusual 8(14)-unsaturation found in the sterols produced by aerobic methanotrophs, dinoflagellates, nematodes, and marine sponges, remains unknown. Here, we utilize a heterologous expression system to identify a bacterial 8,14-sterol reductase (8,14-Bsr) responsible for generating the 8(14)-unsaturation in the aerobic methanotroph *Methylococcus capsulatus*. This enzyme modifies the direct product of C-14 demethylation, reducing one double bond in the nuclear core structure and isomerizing the other to produce an 8(14)-sterol. We subsequently tested the requirement of putative active site residues for catalysis through site directed mutagenesis, identifying residues likely involved in interacting with the sterol substrate and directly catalyzing this reaction. Bioinformatic analysis of the distribution of 8,14-Bsr reveals it is unique to the bacterial domain, found primarily in the *Methylococcaceae* family, the Mycobacteriales order, and yet uncultured members of the Myxococcota phylum. Further phylogenetic analysis of 8,14-Bsr suggests it shares an evolutionary history with the C-14 demethylase in these organisms and that these two enzymes were likely inherited together. These results provide insight into novel sterol biochemistry, further delimiting sterol biosynthesis in the bacterial domain from eukaryotes and illustrating the importance of molecular characterization to identify bacterial proteins that interact with sterols.

**Importance:** Many of the eukaryotic proteins required for the biosynthesis of sterols, such as cholesterol, have been characterized. However, the pathways governing analogous processes in the bacterial domain are less characterized. Here, we identify an 8,14-sterol reductase in aerobic methanotrophs. This enzyme carries out a unique biochemical reaction, saturating and isomerizing double bonds in the nuclear core structure to produce an 8(14)-sterol. This reductase is restricted to the bacterial domain, further separating the evolution of bacterial sterol production from eukaryotes. Additionally, we find this reductase is prevalent in members of the sterol degrading order Mycobacteriales, highlighting a potential role for this protein in the remodeling of host sterol production by these pathogens.

## Introduction

Sterols, such as cholesterol, have long been characterized in a eukaryotic context, with a rich body of literature describing the biosynthesis, regulation, and physiology of these lipids in a variety of eukaryotes (1–5). Sterols are also produced and modified sporadically throughout the bacterial domain (6–9). However, much less is known about the biology of these lipids in bacteria compared to their eukaryotic counterparts. Among bacterial sterol synthesizers, aerobic methanotrophs from the family *Methylococcaceae* have proved a tractable system from which to explore bacterial sterol production. Studies focused on these bacteria have revealed novel proteins involved in both the biosynthesis and trafficking of these lipids, setting bacterial sterol production apart from analogous processes in eukaryotes (10, 11). Such discoveries have challenged assumptions about the eukaryotic origins of sterol biosynthesis in bacteria and encouraged a more nuanced discussion about the early evolution of this ancient biosynthetic pathway. Yet there are several steps in bacterial sterol biosynthesis that remain unexplored and characterization of these missing biosynthetic steps could provide further insight into the evolution of sterol biosynthesis.

The sterols produced by *Methylococcaceae* are characterized by an unusual 8(14)-unsaturation in the core ring structure (6, 12). The distinctive structure of these sterols, alongside their unique isotopic signature derived from methanotrophy, has allowed these lipids to serve as biomarkers, or molecular fossils, for aerobic methanotrophy in both modern and ancient environments (13–15). Sterols with an 8(14)-unsaturation can also be found in other members of both the eukaryotic and bacterial domains. Lipids with this modification are the primary sterol produced by toxic dinoflagellate species belonging to the family *Kareniaceae* (16–18). The biophysical properties of these lipids have been proposed as a key mechanism for immunity to the toxins produced by *Kareniaceae* in both the dinoflagellates themselves and parasites specific to these organisms (19). 8(14)-Unsaturations also characterize the hydroxylated sterols found in sponges, corals, fungi, and a myxobacterium (20–24). While the capacity to produce an 8(14)-unsaturation occurs in both eukaryotes and bacteria, the proteins responsible for carrying out this step have yet to be characterized in either domain. Additionally, most aerobic methanotrophs lack the canonical C-14 reductase required to saturate the Δ14 double bond produced by C-14 demethylation. It remains unclear if the Δ14 double bond in these bacteria is saturated in a distinct step from production of the 8(14)-unsaturation, as is seen in eukaryotic sterol biosynthesis pathways, or if these two steps happen simultaneously.

In this study, we use a heterologous expression system to identify a reductase responsible for generating the 8(14)-unsaturation in aerobic methanotrophs. We find that this enzyme takes the product of C-14 demethylation, which has two double bonds in the core ring structure, and both saturates and isomerizes these double bonds to produce an 8(14)-sterol. Through site directed mutagenesis, we then test the role of conserved active site residues, identifying several residues essential for catalysis, and begin to consider the potential mechanism underlying this reaction. We explore the taxonomic distribution of this protein and find this reductase is restricted to the bacterial domain, occurring in other aerobic methanotrophs, uncultured myxobacteria, and members of Mycobacteriales, including the pathogen *Mycobacterium tuberculosis*. Through our phylogenetic analysis we reveal a close evolutionary relationship between this reductase and the C-14 demethylase (Cyp51), which directly precedes it in sterol biosynthesis. Altogether, our findings assign a biochemical function to a previously uncharacterized hypothetical protein found both in environmental sterol producing and pathogenic sterol degrading bacteria while continuing to differentiate bacterial sterol biology from similar processes in eukaryotes.

## Results

### Identification of a bacterial 8,14-sterol reductase

During sterol biosynthesis aerobic methanotrophs, like eukaryotes, demethylate sterols at C-14 via the C-14 demethylase (Cyp51), resulting in a Δ8,14 unsaturation (25). This product is then modified to produce the 8(14)-unsaturation characteristic of the sterols synthesized by this family. However, apart from *Methylomicrobium* species (26), these methanotrophs do not harbor a homolog to the canonical C-14 sterol reductase required to form the Δ8 unsaturation common in eukaryotes, and the enzyme responsible for generating the 8(14)-unsaturation remains unknown (Figure 1). To identify potential candidates for these modifications in aerobic methanotrophs, we used a comparative genomics approach. We searched for genes in *Methylococcus capsulatus* that are also present in *Methylobacter luteus, Methylomonas lenta*, and *Methyloprofundus sedimenti,* members of *Methylococcaceae* that produce sterols, and are absent in the genomes of *Crenothrix polyspora*, *Methylohalobius crimeensis*, and *Methylomonas methanica,* which belong to the same order but do not produce sterols. This approach generated a list of 10 gene candidates (Table S1). From this list, we identified a hypothetical protein that co-localizes with known sterol biosynthesis genes in the genomes of several methanotrophs (Figure 1). This protein does not have sequence homology to any characterized protein but belongs to the nuclear transport factor 2 (NTF2) superfamily, which includes other proteins involved in sterol modification (27). Additionally, this gene is present in the genomes of all sterol-producing aerobic methanotrophs. Given its wide distribution and its co-localization with known sterol biosynthesis genes, we hypothesized that this protein may be involved in performing either the reduction of the Δ14 unsaturation or generation of the 8(14)-unsaturation.

**Figure 1.**
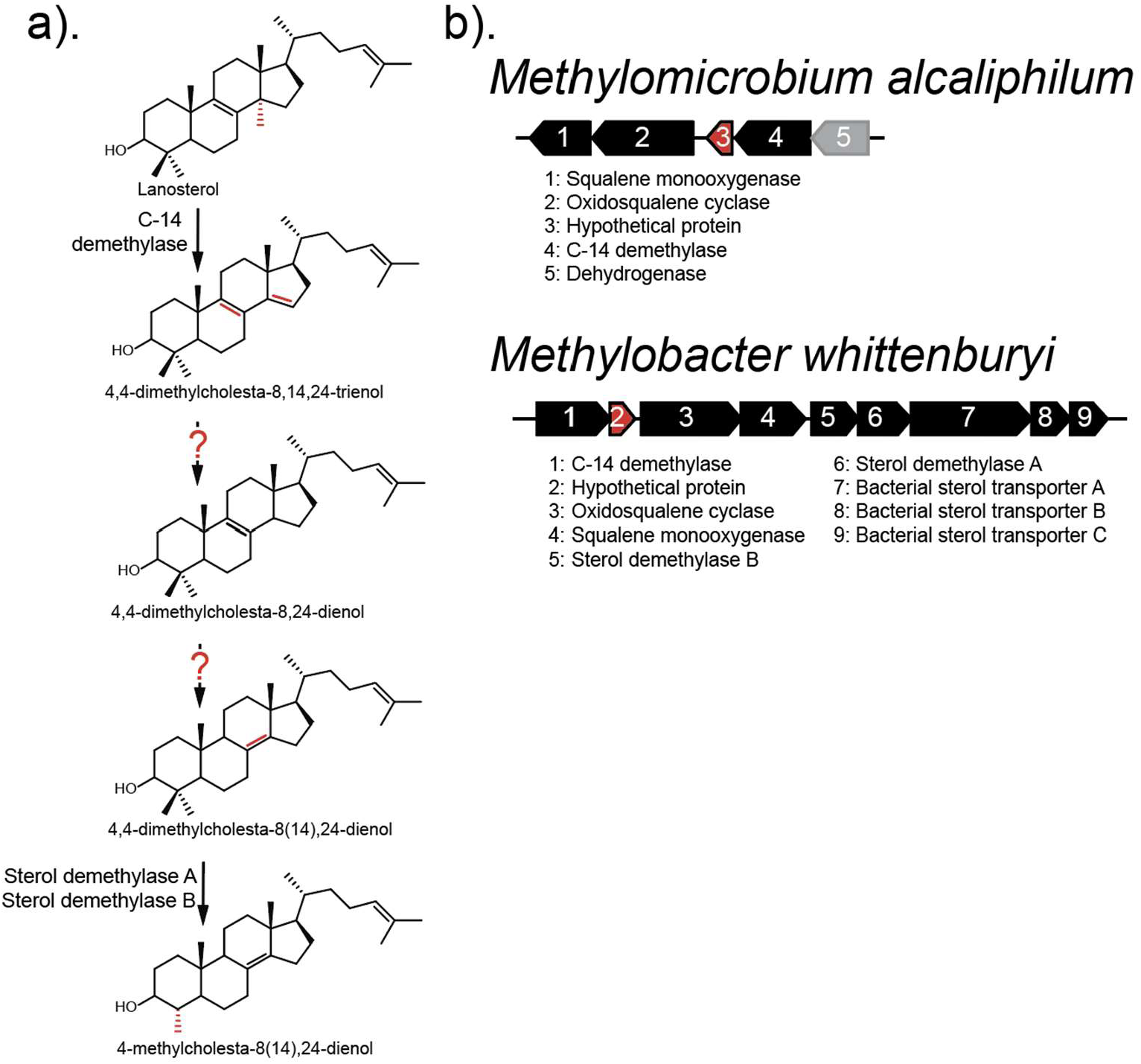
Aerobic methanotrophs produce 8(14)-unsaturated sterols by an unknown mechanism. a) The sterol biosynthesis pathway in aerobic methanotrophs. After cyclization of oxidosqualene, the C-14 methyl group is removed from lanosterol by the C-14 demethylase (Cyp51), generating a Δ14,15 double bond. It is unknown how the Δ14,15 double bond is removed, or the 8(14)-double bond generated in these bacteria. In some methanotrophs, sterols are further demethylated once at C-4 by the sterol demethylase AB (sdmAB) proteins. b) Examples of sterol biosynthesis gene clusters in methanotrophs, featuring known sterol biosynthesis and transport genes (black), a gene without a known function in sterol biosynthesis (grey), and the hypothetical protein characterized in this study (red).

We initially attempted to delete this gene (locus tag: H156DRAFT_MCAT2205) in *Methylococcus capsulatus* through both an unmarked and marked approach, however neither of these strategies were successful (Figure S1). Instead, to test the biochemical function of this protein, we used a heterologous expression system that leverages a series of compatible plasmids to produce the sterol biosynthesis intermediate lanosterol in *Escherichia coli*, which does not produce sterols (28). Expression of the candidate protein MCAT2205 from *M. capsulatus* with lanosterol as a substrate in *E. coli* resulted in no detectable activity, suggesting lanosterol may not be a suitable substrate for this protein. To expand the potential substrates produced by our heterologous expression system, we then introduced the fused C-14 demethylase-ferredoxin homolog (Cyp51) from *M. capsulatus* (locus tag: H156DRAFT_MCAT1746). Expression of the C-14 demethylase resulted in production of 4,4-dimethylcholesta-8,14,24-trienol, a C-14 demethylated sterol with an unsaturation at both Δ14 and Δ8. We then co-expressed the candidate MCAT2205 protein with the C-14 demethylase, and this resulted in complete conversion of all detectable 4,4-dimethylcholesta-8,14,24-trienol to 4,4-dimethylcholesta-8(14), 24-dienol, a product of sterol biosynthesis in aerobic methanotrophs (Figure 2; Figure S2) (29). We further confirmed the production of the dienol sterol by NMR analysis (Figure S3). These results demonstrate that this enzyme is a bacterial 8,14-sterol reductase (8,14-Bsr), and it is sufficient to perform both a reduction and isomerization reaction to remove the Δ14 and Δ8 double bonds, generating the 8(14)-double bond typically observed in the sterols produced by aerobic methanotrophs.

**Figure 2.**
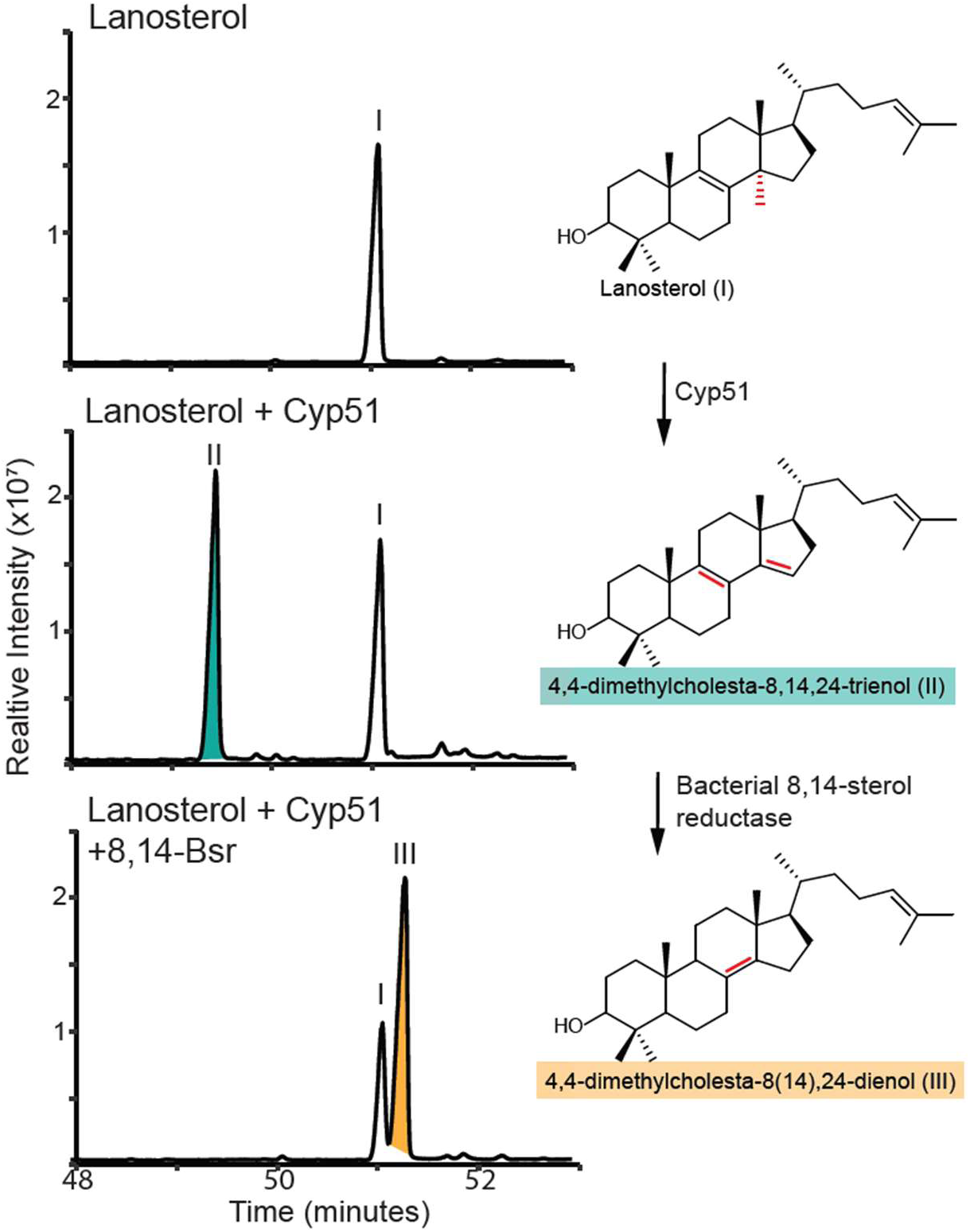
Bacterial 8,14-sterol reductase (8,14-Bsr) generates an 8(14)-unsaturation from 4,4-dimethylcholesta-8,14,24-trienol. Total ion chromatograms from heterologous *E. coli* expression cultures. Expression of the C-14 demethylase (Cyp51) in a lanosterol producing strain of *E. coli* results in production of 4,4-dimethylcholesta-8,14,24-trieneol (II; teal). Co-expression of 8,14-Bsr with Cyp51 results in production of 4,4-dimethylcholesta-8(14), 24-dienol (III; orange). All lipids were derivatized to trimethylsilyl ether groups. Mass spectra of identified sterols are shown in Figure S2.

### Site directed mutagenesis of conserved residues does not separate the reduction and isomerization functions of 8,14-Bsr

Sterol biosynthesis is characterized by examples of bifunctional enzymes, processive reactions, and allylic mechanisms (30–33). We were curious if we could provide further insight into the potential mechanism of 8,14-Bsr, which appears to reduce one double bond in the ring structure and isomerize the other, through site directed mutagenesis of potential active site residues. We hypothesized that if the reaction performed by 8,14-Bsr involves a distinct reduction reaction followed by the isomerization of the remaining double bond, we might be able to mutagenize the active site to result only in the production of a reduced intermediate. To identify potential amino acid residues involved in catalysis, we first compared a structural model of 8,14-Bsr to the structure of ketosteroid isomerase (PBD#1OH0), a well characterized NTF2 family protein that functions on steroids (27, 34). Ketosteroid isomerase (Ksi) harbors a hydrophobic active site pocket found between the alpha helices B and C and catalysis by Ksi requires several tyrosine and aspartic acid residues found within this pocket (35). A structural model of 8,14-Bsr shares the same cone shaped fold as other NTF2 family proteins and, like ketosteroid isomerase, has a hydrophobic pocket between helices B and C. However, the active site residues found in Ksi do not align with those in 8,14-Bsr. Instead, we identified two aspartic acids (D133 and D136) and four tyrosine residues (Y37, Y43, Y78 and Y80) present in this pocket that are largely conserved among other homologs of this protein (Figure S4).

To test if these residues are involved in catalysis, we first individually mutated each aspartic acid residue to alanine and co-expressed these variants with the C-14 demethylase in our *E. coli* heterologous expression system. Mutating the aspartic acid residue at position 133 to alanine (D133A) resulted in the accumulation of the substrate 4,4-dimethylcholesta-8,14,24-trienol, indicating a decrease in 8,14-Bsr function and suggesting this residue is involved in the enzymes mechanism. Conversely, mutating the aspartic acid at position 136 to alanine (D136A) had no effect on enzyme function (Figure 3a). We next individually mutated each tyrosine residue to either phenylalanine, removing the hydroxyl group but maintaining the aromaticity of the side chain, or alanine, removing both the hydroxyl group and aromaticity of the side chain, and co-expressed these variants as well. Mutating the tyrosine at position 80 to either phenylalanine (Y80F) or alanine (Y80A) had no effect on the enzyme function in this system (Figure 3b). The Y37F and Y78F mutations also had no effect on enzyme function. However, mutating these residues to alanine (Y37A or Y78A) resulted in reduced enzyme function, suggesting the aromaticity of these two tyrosine residues plays an important role in performing this reaction, perhaps through pi-stacking interactions with the rings of the sterol substrate (Figure 3c). A broader search of 8,14-Bsr homologs identifies one in the Actinobacterium *Haloenchinothrix aidingensis* with a phenylalanine residue at position 37, illustrating a biologically relevant example of this mutation. Finally, mutating Y43 to either phenylalanine (Y43F) or alanine (Y43A) resulted in decreased enzyme function, suggesting the hydroxyl group of this tyrosine residue may be directly involved in catalysis (Figure 3d).

**Figure 3.**
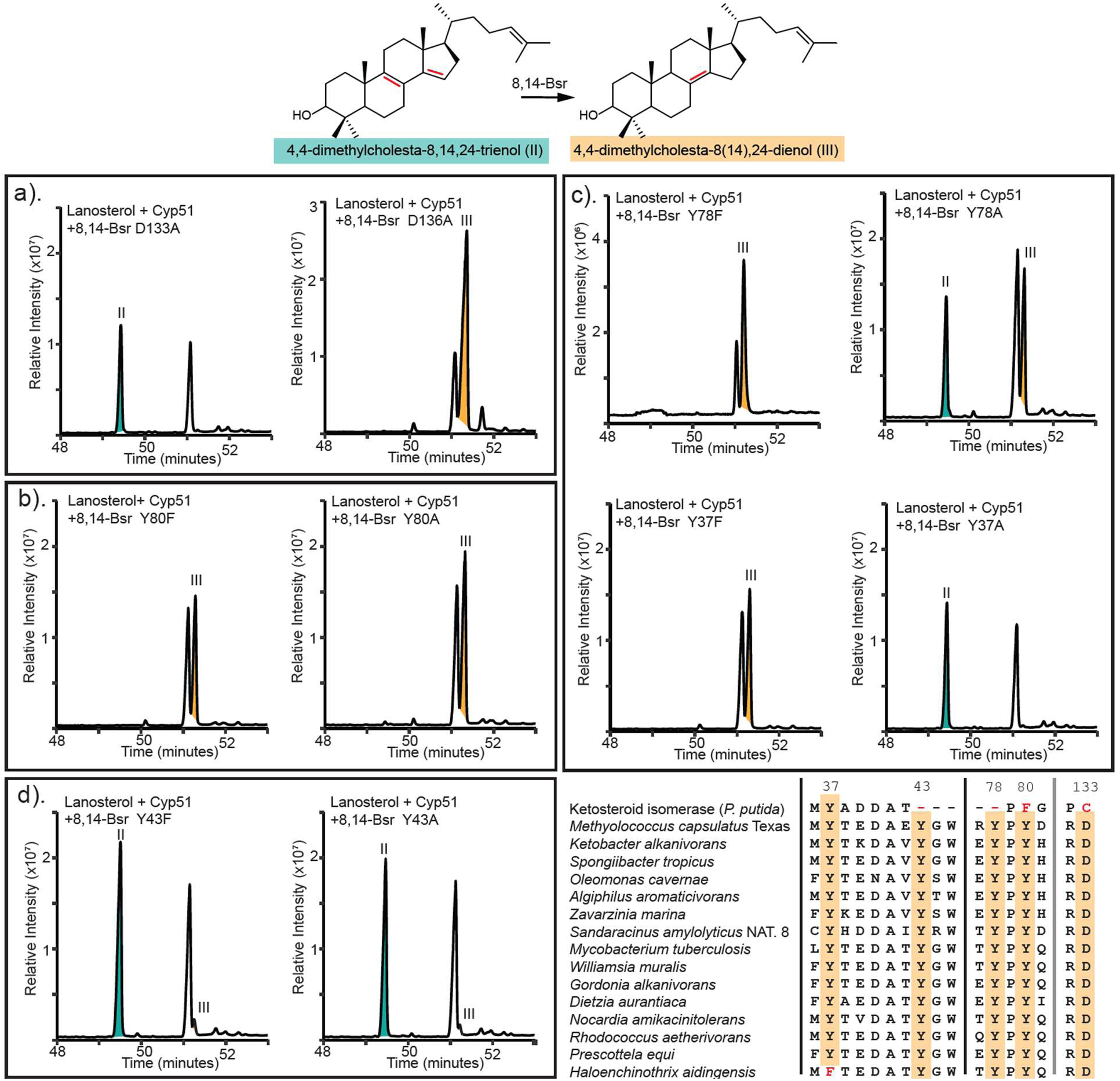
Site directed mutagenesis of conserved residues in 8,14-Bsr reveals residues essential for catalysis. Total ion chromatograms of lipids extracted from heterologous expression of Cyp51 with putative 8,14-Bsr active site pocket mutants. Peaks corresponding to the substrate for 8,14-Bsr, 4,4-dimethylcholesta-8,14,24-trienol, are labeled teal and the product, 4,4-dimethylcholesta-8(14)-dienol, are labeled orange. a) Mutation of the aspartic acid at position 133 and 136 to alanine. b) Mutations of the tyrosine at position 80 to either phenylalanine or alanine, showing no impact on 8,14-Bsr function. c) Mutations of the tyrosine at position 78 and 37 to either phenylalanine or alanine, where only mutations to alanine impact function. d) Mutations of the tyrosine at position 43 to either phenylalanine or alanine, where both mutation to phenylalanine and alanine impact function. An alignment of these conserved residues in 8,14-Bsr homologs is included for reference.

While we were able to identify residues in the active site pocket important for the reaction carried out by 8,14-Bsr, we were unable to generate a mutant which only performs the reduction reaction suggesting that these two steps may be carried out through a single reaction mechanism. This could occur through allylic rearrangement after the reduction of the Δ14 double bond. The canonical Δ14 reductase saturates the Δ14 double by the addition of a hydride from NADPH at C-15 followed by protonation at C-14 to generate a Δ8 sterol (36). We propose that 8,14-Bsr, like the canonical Δ14 reductase, reduces its substrate by hydride addition at C-15. It is possible this enzyme also uses NADPH as a hydride donor during this process, however 8,14-Bsr has no sequence homology to known cofactor binding sites. Hydride addition is then followed by protonation of an allylic anion intermediate at C-9 to generate an unsaturation at 8(14) (Figure S5). Additional biochemical and structural characterization of this enzyme is needed to confirm this mechanism, identify the role of potential cofactors, and determine the function of specific residues in catalyzing this unique reaction.

### 8,14-Bsr is a sterol biosynthesis protein restricted to the bacterial domain

Sterols with an 8(14)-unsaturation are found in a diverse array of bacteria and eukaryotes. To identify if 8,14-Bsr is responsible for 8(14)-double bond formation in these other organisms, we conducted BLASTp and tBLASTn searches against the genomes and transcriptomes present in the IMG and NCBI databases (e-value cutoff e-05). Among eukaryotes, including those that produce 8(14)-sterols such as dinoflagellates and nematodes (37, 38), we did not identify any homologs to the bacterial 8,14-sterol reductase. Furthermore, many of the eukaryotes that produce 8(14)-sterols harbor homologs to the C-14 reductase (Table S2), suggesting that these C-14 reductase homologs also protonate sterols at C-9 during reduction or that these eukaryotes might require an additional, unidentified enzyme to generate the 8(14)-unsaturation. In either case, the biosynthesis of 8(14)-sterols provides an additional instance of independent evolution between sterol production in the eukaryotic and bacterial domains.

While 8,14-Bsr is absent from the eukaryotic domain, homologs are present in a small but diverse group of bacteria. This includes all sterol-producing aerobic methanotrophs, even *Methylomicrobium* species which also harbor a C-14 reductase. 8,14-Bsr homologs from *Methylococcaceae* cluster together and broadly recapitulate 16S rRNA phylogeny (Figure S6). These phylogenetic patterns, alongside the wide distribution of the bacterial 8,14-sterol reductase in aerobic methanotrophs, suggests that the 8(14)-double bond is an ancestral feature of sterol biosynthesis in the family *Methylococcaceae*. 8,14-Bsr homologs are also found in the genomes of γ-proteobacteria, α-proteobacteria, Mycobacteriales, and uncultured myxobacteria. Several of these bacteria are characterized sterol degraders and consume sterols acquired from the environment as a carbon source (39–42). To confirm these homologs can function as 8(14)-sterol reductases, we co-expressed homologs from the uncultured myxobacteria *Sandaracinus amylolyticus* NAT-131 and the sterol degrader *Mycobacterium tuberculosis* in our *E. coli* heterologous system (Figure 4a). In both cases, these homologs produced the same 8(14)-sterols we observe from heterologous expression of the *M. capsulatus* homolog, suggesting the capacity to synthesize 8(14)-sterols, either de novo or from exogenously acquired sterol precursors, is present in these disparate phyla as well.

**Figure 4.**
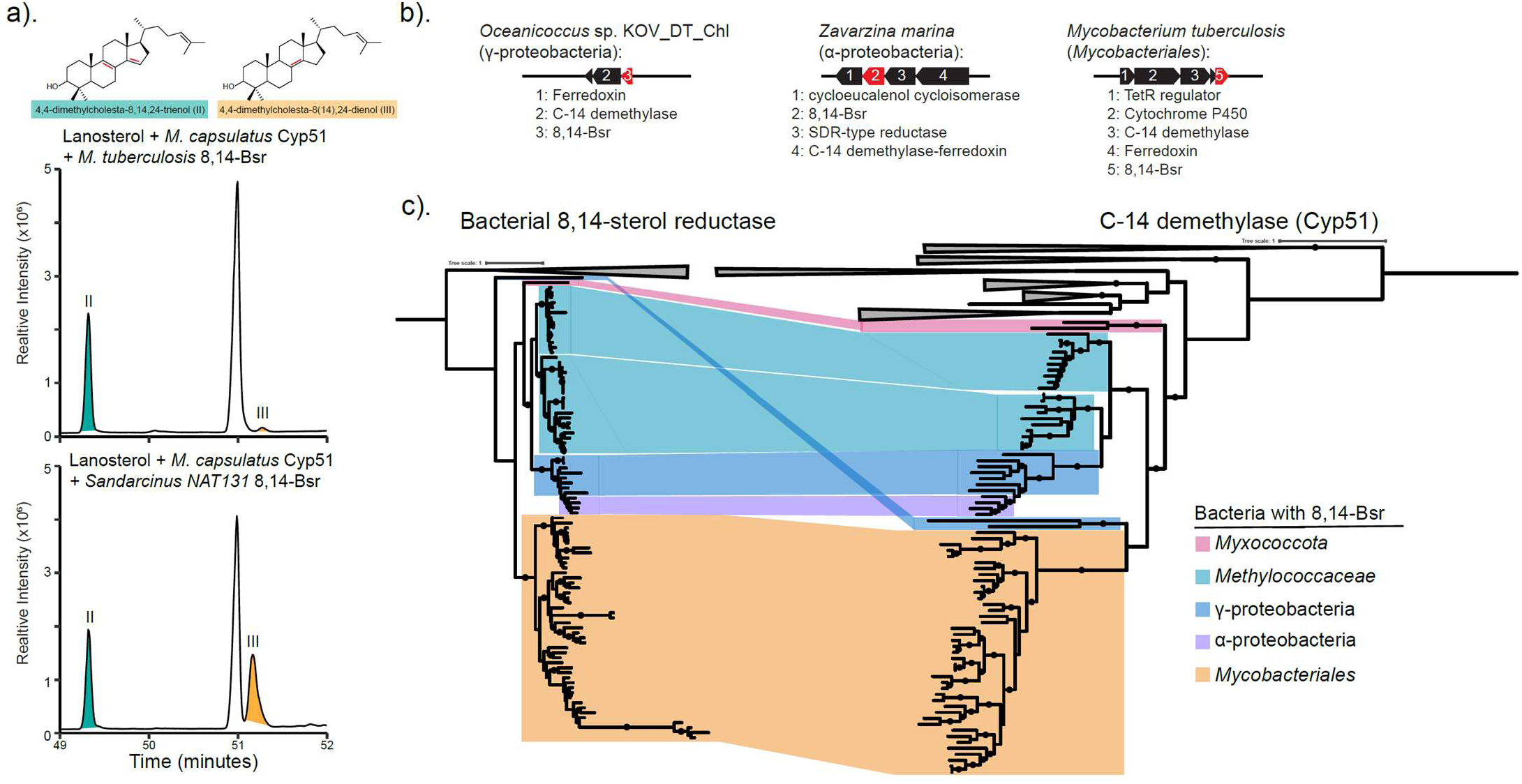
The bacterial 8,14-sterol reductase (8,14-Bsr) is found in diverse sterol producing and sterol degrading bacteria. a) Total ion chromatograms of lipids extracted from heterologous expression of Cyp51 from *Methylococcus capsulatus* Texas with either the 8,14-Bsr homolog from *Mycobacterium tuberculosis* or the uncultured myxobacteria *Sandaracinus* sp. NAT131. Peaks corresponding to the substrate for 8,14-Bsr, 4,4-dimethylcholesta-8,14,24-trienol, are labeled teal (II) and the product, 4,4-dimethylcholesta-8(14)-dienol, orange (III). b) Representative gene clusters from diverse bacteria where Cyp51, the Cyp51 ferredoxin, and 8,14-Bsr are co-localized. 8,14-Bsr is featured in red. c) Maximum likelihood phylogenetic trees of bacterial 8,14-sterol reductase (8,14-Bsr) with ketosteroid isomerase (Ksi) as an outgroup and Cyp51 with Cyp4 as an outgroup were generated using IQTree with the model of best fit and 5000 ultrafast bootstrap replicates. Branches with bootstrap support values >90 are denoted with a black circle. Sequences from different taxonomic groups which harbor 8,14-Bsr homologs are mapped onto the Cyp51 tree and colored by their taxonomic group as follows: Myxococcota (pink), *Methylococcaceae* (Cyan), γ-proteobacteria (dark blue), α-proteobacteria (purple), and Mycobacteriales (orange).

Across these bacteria, 8,14-Bsr often co-localizes with the C-14 demethylase (Figure 4b). Phylogenetic analysis of the C-14 demethylases that co-occur with 8,14-Bsr reveals that these C-14 demethylase homologs form a distinct clade from C-14 demethylases found in other bacteria and eukaryotes, even in the case of sequences from Mycobacteriales and *Streptomyces* despite both belonging to the phylum Actinomycetes (Figure 4c; Figure S7-S8). Additionally, these C-14 demethylase homologs are all fused or co-localized with their cognate ferredoxin (26), which is not observed of other C-14 demethylases (Figure S9). Broadly, the branching pattern of this clade of C-14 demethylases largely mirrors what we observe from 8,14-Bsr, suggesting these two enzymes have co-evolved in the sterol-producing and sterol-degrading bacteria that harbor this distinct protein pair.

## Discussion

Here, we identified a sterol reductase that generates an unsaturation at 8(14) through the reduction and allylic rearrangement of the Δ14 and Δ8 double bonds to result in the unusual nuclear core structure configuration that characterizes sterols in aerobic methanotrophs. The bacterial 8,14-sterol reductase shares no sequence homology to any proteins with known functions, illustrating the importance of coupling molecular characterization to bioinformatic analysis when identifying bacterial proteins involved in sterol biosynthesis, as bioinformatic analyses alone were not sufficient to predict the function of this enzyme. While many proteins and motifs that are diagnostic for sterol binding have been identified in eukaryotes, they are largely absent in bacterial sterol producers. Instead, sterol-producing bacteria continue to present an opportunity to uncover novel sterol biochemistry, better informing how we consider the biological and evolutionary relationships governing sterol modification across the bacterial domain. This has been particularly true in aerobic methanotrophs, which have been previously shown to use unique proteins in both the synthesis and transport of sterols (10, 11). By continuing to characterize bacterial sterol biosynthesis and physiology, we will gain a more complete picture of the biochemistry surrounding these important lipids.

Our work also illustrates the biotechnological utility in developing heterologous systems to study complex biosynthesis pathways. Expression of the fused C-14 demethylase-ferredoxin homolog from *M. capsulatus* resulted in production of the fertility hormone follicular fluid-meiosis activating sterol (FF-MAS). FF-MAS is found in high abundance in mammalian follicular fluid where it is involved in regulation of oocyte maturation (43, 44) and this accumulation is important in the resumption of meiosis in human, mouse, and porcine oocytes during ovulation (45–47). While FF-MAS has proved promising in increasing successful culturing of porcine and mouse oocytes in vitro (47–50), it remains unclear if there may be a role for FF-MAS in the culturing of human oocytes for in vitro fertilization treatments (51, 52). Complicating the study and use of this hormone in a clinical setting, FF-MAS is not present in high concentrations from extractable sources. Currently, a 9-step chemical synthesis scheme for FF-MAS has been developed for production (53). Heterologous expression may present an alternative and more sustainable pathway for generating FF-MAS and, with further optimization, prove a useful source of this lipid for both clinical and research purposes.

Here, we also show that 8,14-Bsr is entirely absent from the eukaryotic domain. The absence of this protein from eukaryotic genomes, particularly those of the dinoflagellates, sponges, and nematodes that produce 8(14)-unsaturated sterols, points to convergent evolution underlying this modification in the two domains. However, how eukaryotes generate 8(14)-sterols remains an open question. This reaction could conceivably be catalyzed by an ortholog to the canonical C-14 reductase that protonates at the C-9 position, as we have proposed for 8,14-Bsr, to generate an 8(14)-sterol. Alternatively, generation of the 8(14)-unsaturation could also be catalyzed by an isomerase after the canonical C-14 reductase produces a Δ8 sterol. The treatment of *Karenia brevis*, a toxic dinoflagellate that synthesizes 8(14)-sterols, with the fungicide fenpropidine, an inhibitor of both the C-14 reductase and C-8 isomerase, resulted in accumulation of Δ8,14,22 sterols (16), suggesting an ortholog to one of these two enzymes is responsible for generating 8(14)-sterols. Additionally, *C. elegans*, which modifies exogenously acquired sterols, has been hypothesized to generate 8(14) sterols through an ortholog to the eukaryotic C-8 sterol isomerase that would isomerize the Δ7 double bond to 8(14) (54). Further functional characterization of both the C-14 reductase and C-8 isomerase orthologs in these organisms could resolve these outstanding questions, better clarifying the evolutionary trajectory of 8(14)-sterol production in eukaryotes.

The independent evolution of enzymes responsible for producing 8(14)-unsaturated sterols in eukaryotes and bacteria also highlights the biological importance of these small modifications to the biophysical properties and physiological functions of these lipids in the cell. In the case of either domain, the physiological role of the 8(14)-unsaturation remains unknown. Sterols with an 8(14)-unsaturation from the toxic dinoflagellate *K. brevis* have a protective effect against the toxins produced by this organism (19). These sterols have also been proposed as a potential deterrent for predation as they might interfere with predator sterol metabolism (55). However, the sterols from these dinoflagellate species are also characterized by unique modifications to the sterol side chain (38) and the effect of these side chain modifications versus the nuclear core unsaturation on membrane dynamics is unknown. Similarly, we do not know the physiological function of this modification in sterol-producing bacteria, such as the aerobic methanotrophs. Yet it is interesting to note that while 8,14-Bsr is present in all sterol producing aerobic methanotrophs, *Methylomicrobium* species also harbor a C-14 reductase and C-8 isomerase suggesting these bacteria are capable of producing sterols with different double bond configurations in the core structure. This pathway bifurcation again highlights the potential importance of these small sterol modifications in *Methylococcaceae* and suggests a possibility for specific functions tied to specific sterols.

Finally, our confirmation of 8,14-Bsr activity in sterol-degrading bacteria also raises questions about the function of these enzymes in Mycobacteriales. Pathogens from this family are well known for their ability to catabolize host sterols, specifically cholesterol, and the enzymes involved in this process are essential for pathogenicity (56). However, less is known about how these bacteria interact with other sterols produced by the host. The C-14 demethylase from several *Mycobacterium* species have been crystalized and characterized, demonstrating that these enzymes are functional C-14 demethylases (57–59). While neither the C-14 demethylase nor 8,14-Bsr are essential to cell viability or pathogenicity (60–62), the presence of these two genes in *Mycobacterium* species would suggest these host-associated bacteria have the capacity to modify exogenously acquired lanosterol. Indeed, a study found that when fed lanosterol as a carbon source, a *Mycobacterium* sp. converted this sterol into a C_19_ steroid with an 8(14)-unsaturation in the ring structure (63, 64). Mycobacteria include intracellular pathogens that persist in macrophages. Macrophages accumulate lanosterol early in bacterial infection in response to Toll-like receptor 4 activation, triggering downstream signaling pathways and increasing membrane fluidity to allow for increased phagocytosis (65). Perhaps the C-14 demethylase and 8,14-Bsr are used by *Mycobacterium* to remodel sterol production in these macrophages during infection. In support of this, 8,14-Bsr was found to be upregulated in *M. tuberculosis* early in macrophage infection (66, 67). Further research investigating the role of lanosterol degradation in *Mycobacterium* physiology could yield additional insight into the pathogenic life cycle of these bacteria. Altogether, our findings provide a foundation for probing new questions around sterol physiology in both sterol-producing and sterol-degrading bacteria while highlighting the utility in studying environmental bacteria, such as *Methylococcaceae*, to explore basic and fundamental biological processes.

## Methods

### Bacterial Culture

For genomic DNA extraction, *Methylococcus capsulatus* Texas was grown in liquid culture on NMS media supplemented with methane for 3 days at 37°C (68). *Escherichia coli* DH10B was used for all cloning and expression experiments and cultured on either Luria Broth (LB) or TYGPN media. In heterologous expression experiments, *E. coli* DH10B was cultured in TYPGPN medium, at an initial temperature of 37°C and cultures were shifted to 30°C at expression induction for 40 hours, shaking at 220 rpm. Cultures were supplemented with 100µg/ml of carbenicillin, 20µg/ml of chloramphenicol, 30µg/ml of kanamycin, and/or 15µg/ml of gentamycin as needed

### Molecular Cloning

All oligonucleotides and plasmids used in this study are described in Table S3 and S4. Oligonucleotide primers were ordered from Integrated DNA Technologies (Coralville, IA). Plasmids were isolated from *E. coli* through the GeneJET Plasmid Miniprep Kit (Thermo Scientific). DNA fragments used in cloning were gel purified through the GeneJET Gel Extraction Kit (Thermo Scientific). Constructed plasmids were sequenced by ELIM Biopharm (Hayward, CA).

All plasmids were constructed using sequence and ligation independent cloning (SLIC) (69). Briefly, gene fragments were amplified using Phusion DNA Polymerase (New England Biolabs) with complementary plasmid insertion site overhangs and gel purified. The expression plasmids pBAD1031K or pSRKGm-lacUV5-rbs5 were linearized by digestion using either NcoI or NdeI (New England Biolabs), respectively. Complementary overhangs were created on both gel-purified gene fragments and linearized plasmids by incubation with T4 DNA Polymerase (MilliporeSigma) without nucleotides. Vectors and gene fragments were then annealed and transformed by electroporation without ligation into *E. coli*.

### Site Directed Mutagenesis

Site directed mutagenesis was performed by PCR amplification of the vector with a single oligonucleotide with the desired change. This reaction was performed using 2.5 units of PfuUltra II Fusion HS DNA Polymerase (Agilent) with the oligonucleotide (0.2 μM), 0.2 mM dNTPs, and 50ng plasmid DNA in 1x PfuUltra buffer and a 1-minute per kilobase pair extension time at 68 °C. The reaction was then digested with DpnI (New England Biolabs) and transformed by electroporation into *E. coli*. The mutation was verified through plasmid sequencing by ELIM Biopharm (Hayward, CA).

### Heterologous Expression

Biosynthesis genes for production and cyclization of oxidosqualene were expressed from a set of compatible plasmids in *E. coli* DH10B (29). Heterologous expression strains were constructed as described in Table S5. Genes of interest were expressed from either the IPTG-inducible plasmid pSRKGm-*lac*UV5-rbs5 or the arabinose-inducible plasmid pBAD1031K. Strains were cultured in 20ml of TYPGN media supplemented with the necessary antibiotics, at 37°C, shaking at 220 rpm, until mid-log phase. At mid-log phase, expression was induced with 500µM IPTG and 0.2% (wt: vol) arabinose and cultures were shifted to 30°C, shaking at 220 rpm, for 30-40 hours before harvesting cultures by centrifugation at 4,500xg for 10 minutes.

### Lipid Extraction

Lipids were extracted through a modified Bligh-Dyer extraction (70). Cell pellets were sonicated in a water bath in 10:5:4 methanol: dichloromethane (DCM): water (vol: vol: vol) for 1 hour. Lipids were then phase separated by addition of two times the volume 1:1 DCM: water (vol: vol) and the organic layer removed and dried under nitrogen gas. Lipids were then derivatized to either trimethylsilyl ethers by adding 1:1 (vol: vol) Bis(trimethylsilyl)trifluoroacetamide: pyridine and heating at 70°C for 1 hour or derivatized to acetyl groups by adding 1:1 (vol: vol) acetic anhydride: pyridine and heating at 70°C for 1 hour and resuspending in DCM.

### Gas Chromatography Mass Spectrometry (GC-MS)

Sterols were separated on an Agilent 7890B Series GC using a 60 m Agilent DB17HT column (60 m x 0.25 mm i.d. x 0.1 μm film thickness) with helium as the carrier gas at constant flow of 1.1 mL/min and programed as follows: 100°C for 2 minutes; then 8°C/minute to 250°C and held for 10 minutes; then 3°C/minutes to 330°C and held for 17 minutes. 2 μL of each sample was injected in splitless mode at 250°C. The GC was coupled to a 5977 A Series MSD with the ion source held at 230°C and operated at 70 eV in EI mode scanning from 50-850 Da in 0.5 s. Lipids were analyzed using Agilent MassHunter Qualitative Analysis (B.06.00) and identified based on retention time and spectra by comparison to previously confirmed laboratory standards, spectra deposited in the National Institute of Standards and Technology (NIST) databases or previously published spectra as well as through NMR analysis.

### NMR Analysis of 4,4-dimethylcholesta-8(14), 24-dienol

The lipids were fractionated by preparative thin-layer chromatography (TLC) on glass-backed plates (10 cm in length) coated with a 0.25 mm layer of silica gel 60 F254 using hexane/EtOAc 4:1 as the developing solvent. The TLC fractions were characterized by NMR using a Bruker Avance III HD with an Ascend 800 MHz magnet and a 5 mm TCI cryoprobe at 30°C using deuterated chloroform (CDCl3) as the solvent. Calibration was by the residual solvent signal (7.26 ppm). NMR spectral data shown in Figure S3.

### Bioinformatics

Homologs of the bacterial 8(14)-sterol reductase (locus tag: H156DRAFT_MCAT2205) and the C-14 demethylase (locus tag: H156DRAFT_MCAT1746) were identified using a BLASTp search (e-value cutoff of e-30) of the cultured genomes available in the JGI IMG and NCBI databases (71–73). BLAST searches against eukaryotic genomes also included a tBLASTn search of the transcriptome database in NCBI (e-value cutoff e-05). Protein alignments were performed in MAFFT (version 7.526) using the L-INS-I algorithm with 1000 iterations (74, 75). Phylogenetic trees were generated using IQ-TREE2 (version 3.0.1). Modelfinder was used to identify the best fit model for both bacterial 8(14)-sterol reductase and the C-14 demethylase trees and a maximum likelihood tree was generated with 5000 ultrafast bootstrap replicates (76, 77). Phylogenetic trees were then edited for figures using the iTOL website (http://itol.embl.de/).

Models of bacterial 8,14-sterol reductase homologs were generated using ColabFold alphafold2 platform (version 1.5.5)(78). Ketosteroid isomerase was identified as a structurally similar protein based on structural similarity search through the RCSB Protein Database using the alpha model of the *M. capsulatus* bacterial 8,14-sterol reductase homolog (79). Bacterial 8,14-sterol reductase was aligned to the crystal structure of ketosteroid isomerase (PDB: 1OH0) in Pymol (version 2.5.7) using the cealign function. The active site pocket of both proteins was visualized using the set_surface_mode, 2 command and colored by hydrophobicity using the color h script (80).

## Acknowledgements

We thank the members of the Welander lab for their helpful discussions and suggestions. Portions of the experimental work were performed in the Stanford Geomicrobiology Shared Laboratories Core Facility (RRID:SCR_025000). Funding for this study was provided to P.V.W. by National Science Foundation Award 1919153. The acquisition of the 800 MHz NMR spectrometer at SUNY-ESF was made possible by NIH grant S10 OD012254.

## Notes

### Competing Interest Statement

The authors have declared no competing interest.

### Summary of Updates

The NMR data that was previously marked as data not shown was added to the supplemental file (Figure S3). A new funder and grant number was added to the acknowledgments in the main text.

## References

1. Bloch K. 1965. The Biological Synthesis of Cholesterol. Science 150:19–28.

2. Róg T, Pasenkiewicz-Gierula M, Vattulainen I, Karttunen M. 2009. Ordering effects of cholesterol and its analogues. Biochimica et Biophysica Acta (BBA) - Biomembranes 1788:97–121.

3. Anderson RH, Sochacki KA, Vuppula H, Scott BL, Bailey EM, Schultz MM, Kerkvliet JG, Taraska JW, Hoppe AD, Francis KR. 2021. Sterols lower energetic barriers of membrane bending and fission necessary for efficient clathrin-mediated endocytosis. Cell Reports 37:110008.

4. Nes WD. 2011. Biosynthesis of Cholesterol and Other Sterols. Chem Rev 111:6423–6451.

5. Eberlé D, Hegarty B, Bossard P, Ferré P, Foufelle F. 2004. SREBP transcription factors: master regulators of lipid homeostasis. Biochimie 86:839–848.

6. Wei JH, Yin X, Welander PV. 2016. Sterol Synthesis in Diverse Bacteria. Front Microbiol 7:990.

7. Pearson A, Budin M, Brocks JJ. 2003. Phylogenetic and biochemical evidence for sterol synthesis in the bacterium Gemmata obscuriglobus. Proceedings of the National Academy of Sciences 100:15352–15357.

8. Kohl W, Gloe A, Reichenbach H. 1983. Steroids from the Myxobacterium Nannocystis exedens. Microbiology 129:1629–1635.

9. Bird CW, Lynch JM, Pirt FJ, Reid WW, Brooks CJW, Middleditch BS. 1971. Steroids and Squalene in Methylococcus capsulatus grown on Methane. Nature 230:473–474.

10. Lee AK, Banta AB, Wei JH, Kiemle DJ, Feng J, Giner J-L, Welander PV. 2018. C-4 sterol demethylation enzymes distinguish bacterial and eukaryotic sterol synthesis. Proceedings of the National Academy of Sciences 115:5884–5889.

11. Zhai L, Bonds AC, Smith CA, Oo H, Chou JC-C, Welander PV, Dassama LM. 2024. Novel sterol binding domains in bacteria. eLife 12:RP90696.

12. Schouten S, Bowman JP, Rijpstra WIC, Sinninghe Damsté JS. 2000. Sterols in a psychrophilic methanotroph, Methylosphaera hansonii. FEMS Microbiology Letters 186:193–195.

13. Bouloubassi I, Nabais E, Pancost RD, Lorre A, Taphanel M-H. 2009. First biomarker evidence for methane oxidation at cold seeps in the Southeast Atlantic (REGAB pockmark). Deep Sea Research Part II: Topical Studies in Oceanography 56:2239–2247.

14. Elvert M, Niemann H. 2008. Occurrence of unusual steroids and hopanoids derived from aerobic methanotrophs at an active marine mud volcano. Organic Geochemistry 39:167–177.

15. Coolen MJL, Hopmans EC, Rijpstra WIC, Muyzer G, Schouten S, Volkman JK, Sinninghe Damsté JS. 2004. Evolution of the methane cycle in Ace Lake (Antarctica) during the Holocene: response of methanogens and methanotrophs to environmental change. Organic Geochemistry 35:1151–1167.

16. Leblond JD, Roche SA, Porter NM, Howard JC, Dunlap NK. 2011. Sterol biosynthesis in the harmful marine dinoflagellate, Karenia brevis: Identification of biosynthetic intermediates produced during exposure to the fungicide fenpropidine. Phycological Research 59:54–63.

17. Leblond JD, Sabir K, Whittemore HL. 2023. Sterol Composition of the Peridinin-Containing Dinoflagellate *Gertia stigmatica*, a Member of the Kareniaceae without a Canonical Haptophyte-Derived Plastid. Protist 174:125939.

18. Mooney BD, Nichols PD, Salas MFD, Hallegraeff GM. 2007. LIPID, FATTY ACID, AND STEROL COMPOSITION OF EIGHT SPECIES OF KARENIACEAE (DINOPHYTA): CHEMOTAXONOMY AND PUTATIVE LIPID PHYCOTOXINS1. Journal of Phycology 43:101–111.

19. Place A, Harvey H, Bai X, Coats D. 2006. Sneaking under the toxin surveillance radar: parasitism and sterol content. African Journal of Marine Science 28:347–351.

20. Santafé G, Paz V, Rodríguez J, Jiménez C. 2002. Novel Cytotoxic Oxygenated C29 Sterols from the Colombian Marine Sponge *Polymastia t enax*. J Nat Prod 65:1161–1164.

21. Sun Y, Tian L, Huang J, Li W, Pei Y. 2006. Cytotoxic sterols from marine-derived fungus Pennicillium sp. Natural Product Research 20:381–384.

22. Ishii T, Matsuura H, Zhaoqi Z, Vairappan CS. 2009. A New 4α-Methylated Sterol from a Nephthea sp. (Nephtheidae) Bornean Soft Coral. 9. Molecules 14:3360–3366.

23. Akihisa T, Tanaka N, Yokota T, Tanno N, Tamura T. 1991. 5α-cholest-8(14)-en-3β-ol and three 24-alkyl-Δ8(14) -sterols from the bulbils of *Dioscorea batatas*. Phytochemistry 30:2369–2372.

24. Gawas D, Garcia R, Huch V, Müller R. 2011. A Highly Conjugated Dihydroxylated C28 Steroid from a Myxobacterium. J Nat Prod 74:1281–1283.

25. Jackson CJ, Lamb DC, Marczylo TH, Warrilow AGS, Manning NJ, Lowe DJ, Kelly DE, Kelly SL. 2002. A Novel Sterol 14α-Demethylase/Ferredoxin Fusion Protein (MCCYP51FX) from Methylococcus capsulatusRepresents a New Class of the Cytochrome P450 Superfamily *. Journal of Biological Chemistry 277:46959–46965.

26. Li X, Roberti R, Blobel G. 2015. Structure of an integral membrane sterol reductase from Methylomicrobium alcaliphilum. Nature 517:104–107.

27. Kim SW, Cha S-S, Cho H-S, Kim J-S, Ha N-C, Cho M-J, Joo S, Kim KK, Choi KY, Oh B-H. 1997. High-Resolution Crystal Structures of Δ5-3-Ketosteroid Isomerase with and without a Reaction Intermediate Analogue. Biochemistry 36:14030–14036.

28. Banta AB, Wei JH, Gill CCC, Giner J-L, Welander PV. 2017. Synthesis of arborane triterpenols by a bacterial oxidosqualene cyclase. Proceedings of the National Academy of Sciences 114:245–250.

29. Bouvier P, Rohmer M, Benveniste P, Ourisson G. 1976. Delta8(14)-steroids in the bacterium Methylococcus capsulatus. Biochem J 159:267–271.

30. Marijanovic Z, Laubner D, Möller G, Gege C, Husen B, Adamski J, Breitling R. 2003. Closing the Gap: Identification of Human 3-Ketosteroid Reductase, the Last Unknown Enzyme of Mammalian Cholesterol Biosynthesis. Mol Endocrinol 17:1715–1725.

31. Tsai P-L, Zhao C, Turner E, Schlieker C. The Lamin B receptor is essential for cholesterol synthesis and perturbed by disease-causing mutations. eLife 5:e16011.

32. McCarty KD, Sullivan ME, Tateishi Y, Hargrove TY, Lepesheva GI, Guengerich FP. 2023. Processive kinetics in the three-step lanosterol 14α-demethylation reaction catalyzed by human cytochrome P450 51A1. Journal of Biological Chemistry 299.

33. Corey EJ, Virgil SC. 1991. An experimental demonstration of the stereochemistry of enzymic cyclization of 2,3-oxidosqualene to the protosterol system, forerunner of lanosterol and cholesterol. J Am Chem Soc 113:4025–4026.

34. Cho H-S, Choi G, Choi KY, Oh B-H. 1998. Crystal Structure and Enzyme Mechanism of Δ5-3-Ketosteroid Isomerase from Pseudomonas testosteroni,. Biochemistry 37:8325–8330.

35. F Y, Jt B, Mm P, T D, As P, Js F, D H. 2020. Assessment of enzyme active site positioning and tests of catalytic mechanisms through X-ray-derived conformational ensembles. Proceedings of the National Academy of Sciences of the United States of America 117.

36. Akhtar M, Alexander K, Boar RB. 1978. Chemical and Enzymic Studies on the Characterization of Intermediates during the Removal of the 14ac-Methyl-Group in Cholesterol Biosynthesis 169.

37. Chitwood DJ, Lusby WR, Lozano R, Thompson MJ, Svoboda JA. 1984. Sterol metabolism in the nematodeCaenorhabditis elegans. Lipids 19:500–506.

38. Leblond JD, Chapman PJ. 2002. A Survey of the Sterol Composition of the Marine Dinoflagellates Karenia Brevis, Karenia Mikimotoi, and Karlodinium Micrum: Distribution of Sterols Within Other Members of the Class Dinophyceae1. Journal of Phycology 38:670–682.

39. García-Fernández E, Frank DJ, Galán B, Kells PM, Podust LM, García JL, Ortiz de Montellano PR. 2013. A highly conserved mycobacterial cholesterol catabolic pathway. Environmental Microbiology 15:2342–2359.

40. Holert J, Cardenas E, Bergstrand LH, Zaikova E, Hahn AS, Hallam SJ, Mohn WW. 2018. Metagenomes Reveal Global Distribution of Bacterial Steroid Catabolism in Natural, Engineered, and Host Environments. mBio 10.1128/mbio.02345-17.

41. Merino E, Barrientos A, Rodríguez J, Naharro G, Luengo JM, Olivera ER. 2013. Isolation of cholesterol- and deoxycholate-degrading bacteria from soil samples: evidence of a common pathway. Appl Microbiol Biotechnol 97:891–904.

42. Pandey AK, Sassetti CM. 2008. Mycobacterial persistence requires the utilization of host cholesterol. Proceedings of the National Academy of Sciences 105:4376–4380.

43. Byskov AG, Andersen CY, Nordholm L, Thogersen H, Guoliang X, Wassmann O, Andersen JV, Guddal E, Roed T. 1995. Chemical structure of sterols that activate oocyte meiosis. Nature 374:559–562.

44. Baltsen M, Byskov AG. 1999. Quantitation of meiosis activating sterols in human follicular fluid using HPLC and photodiode array detection. Biomed Chromatogr 13:382–388.

45. Grøndahl C, Ottesen JL, Lessl M, Faarup P, Murray A, Grønvald FC, Hegele-Hartung C, Ahnfelt-Rønne L. 1998. Meiosis-Activating Sterol Promotes Resumption of Meiosis in Mouse Oocytes Cultured in Vitro in Contrast to Related Oxysterols. Biol Reprod 58:1297–1302.

46. Cavilla JL, Kennedy CR, Baltsen M, Klentzeris LD, Byskov AG, Hartshorne GM. 2001. The effects of meiosis activating sterol on in-vitro maturation and fertilization of human oocytes from stimulated and unstimulated ovaries*. Human Reproduction 16:547–555.

47. Guo R, Wang X, Li Q, Sun X, Zhang J, Hao R. 2020. Follicular fluid meiosis-activating sterol (FF-MAS) promotes meiotic resumption via the MAPK pathway in porcine oocytes. Theriogenology 148:186–193.

48. Coticchio G, Rossi G, Borini A, Grøndahl C, Macchiarelli G, Flamigni C, Fleming S, Cecconi S. 2004. Mouse oocyte meiotic resumption and polar body extrusion in vitro are differentially influenced by FSH, epidermal growth factor and meiosis-activating sterol. Hum Reprod 19:2913–2918.

49. Faerge I, Strejcek F, Laurincik J, Rath D, Niemann H, Schellander K, Rosenkranz C, Hyttel PM, Grøndahl C. 2006. The effect of FF-MAS on porcine cumulus–oocyte complex maturation, fertilization and pronucleus formation in vitro. Zygote 14:189–199.

50. Hegele-Hartung C, Kuhnke J, Lessl M, Grøndahl C, Ottesen J, Beier HM, Eisner S, Eichenlaub-Ritter U. 1999. Nuclear and Cytoplasmic Maturation of Mouse Oocytes After Treatment with Synthetic Meiosis-Activating Sterol In Vitro. Biol Reprod 61:1362–1372.

51. Bergh C, Group for the CIS, Loft A, Group for the CIS, Lundin K, Group for the CIS, Ziebe S, Group for the CIS, Nilsson L, Group for the CIS, Wikland M, Group for the CIS, Gröndahl C, Group for the CIS, Arce J-C, Group for the CIS. 2004. Chromosomal abnormality rate in human pre-embryos derived from in vitro fertilization cycles cultured in the presence of Follicular-Fluid Meiosis Activating Sterol (FF-MAS). Hum Reprod 19:2109–2117.

52. Hossein T, Shabani r, Artimani T, Pilehvari S, Moghimi M, Mehdizadeh M. 2024. Innovative Approaches to Enhancing Oocyte Competence in patients with polycystic ovarian syndrome: A Microfluidic Study with the Impact of FF-MAS on Rescued In Vitro Oocyte Maturation. https://www.researchsquare.com. Retrieved 14 May 2024.

53. Blume T, Guttzeit M, Kuhnke J, Zorn L. 2003. Two syntheses of FF-MAS. Org Lett 5:1837–1839.

54. Zhou W, Fisher PM, Vanderloop BH, Shen Y, Shi H, Maldonado AJ, Leaver DJ, Nes WD. 2020. A nematode sterol C4α-methyltransferase catalyzes a new methylation reaction responsible for sterol diversity [S]. Journal of Lipid Research 61:192–204.

55. Giner J-L, Faraldos JA, Boyer GL. 2003. Novel sterols of the toxic dinoflagellate karenia brevis (dinophyceae): a defensive function for unusual marine sterols?1. Journal of Phycology 39:315–319.

56. Van der Geize R, Yam K, Heuser T, Wilbrink MH, Hara H, Anderton MC, Sim E, Dijkhuizen L, Davies JE, Mohn WW, Eltis LD. 2007. A gene cluster encoding cholesterol catabolism in a soil actinomycete provides insight into Mycobacterium tuberculosis survival in macrophages. Proceedings of the National Academy of Sciences 104:1947–1952.

57. Podust LM, Poulos TL, Waterman MR. 2001. Crystal structure of cytochrome P450 14α-sterol demethylase (CYP51) from Mycobacterium tuberculosis in complex with azole inhibitors. Proceedings of the National Academy of Sciences 98:3068–3073.

58. Jackson CJ, Lamb DC, Marczylo TH, Parker JE, Manning NL, Kelly DE, Kelly SL. 2003. Conservation and cloning of CYP51: a sterol 14α-demethylase from *Mycobacterium smegmatis*. Biochemical and Biophysical Research Communications 301:558–563.

59. Mohamed H, Child SA, Bruning JB, Bell SG. 2022. A comparison of the bacterial CYP51 cytochrome P450 enzymes from *Mycobacterium marinum* and *Mycobacterium tuberculosis*. The Journal of Steroid Biochemistry and Molecular Biology 221:106097.

60. Eshraghisamani R, Mirto AJ, Wang J, Behr MA, Barkema HW, De Buck J. 2022. Identification of essential genes in Mycobacterium avium subsp. paratuberculosis genome for persistence in dairy calves. Front Microbiol 13.

61. Rifat D, Chen L, Kreiswirth BN, Nuermberger EL. 2021. Genome-Wide Essentiality Analysis of Mycobacterium abscessus by Saturated Transposon Mutagenesis and Deep Sequencing. mBio 12:10.1128/mbio.01049-21.

62. DeJesus MA, Gerrick ER, Xu W, Park SW, Long JE, Boutte CC, Rubin EJ, Schnappinger D, Ehrt S, Fortune SM, Sassetti CM, Ioerger TR. 2017. Comprehensive Essentiality Analysis of the Mycobacterium tuberculosis Genome via Saturating Transposon Mutagenesis. mBio 8:10.1128/mbio.02133-16.

63. Lee S-S, Young L-H, Wang KC. 1991. Microbial Transformation of Cycloartenol, 24-Methylenecycloartanol, and Lanosterol, II. Isolation and Characterization of C_19_ Steroids. J Nat Prod 54:178–183.

64. Wang KC, You B-J, Yan J-L, Lee S-S. 1995. Microbial Transformation of Lanosterol Derivatives with Mycobacterium sp. (NRRL B-3805). J Nat Prod 58:1222–1227.

65. Araldi E, Fernández-Fuertes M, Canfrán-Duque A, Tang W, Cline GW, Madrigal-Matute J, Pober JS, Lasunción MA, Wu D, Fernández-Hernando C, Suárez Y. 2017. Lanosterol Modulates TLR4-Mediated Innate Immune Responses in Macrophages. Cell Reports 19:2743–2755.

66. Zimmermann M, Kogadeeva M, Gengenbacher M, McEwen G, Mollenkopf H-J, Zamboni N, Kaufmann SHE, Sauer U. 2017. Integration of Metabolomics and Transcriptomics Reveals a Complex Diet of Mycobacterium tuberculosis during Early Macrophage Infection. mSystems 2:10.1128/msystems.00057-17.

67. Cornejo-Granados F, López-Leal G, Mata-Espinosa DA, Barrios-Payán J, Marquina-Castillo B, Equihua-Medina E, Zatarain-Barrón ZL, Molina-Romero C, Hernández-Pando R, Ochoa-Leyva A. 2021. Targeted RNA-Seq Reveals the M. tuberculosis Transcriptome from an In Vivo Infection Model. Biology 10:848.

68. Whittenbury R, Phillips KC, Wilkinson JF. 1970. Enrichment, Isolation and Some Properties of Methane-utilizing Bacteria. Microbiology 61:205–218.

69. Li MZ, Elledge SJ. 2012. SLIC: A Method for Sequence- and Ligation-Independent Cloning, p. 51–59. In Gene Synthesis. Humana Press.

70. Bligh EG, Dyer WJ. 1959. A RAPID METHOD OF TOTAL LIPID EXTRACTION AND PURIFICATION. Canadian Journal of Biochemistry and Physiology 10.1139/o59-099.

71. Altschul SF, Madden TL, Schäffer AA, Zhang J, Zhang Z, Miller W, Lipman DJ. 1997. Gapped BLAST and PSI-BLAST: a new generation of protein database search programs. Nucleic Acids Research 25:3389–3402.

72. Chen I-MA, Markowitz VM, Palaniappan K, Szeto E, Chu K, Huang J, Ratner A, Pillay M, Hadjithomas M, Huntemann M, Mikhailova N, Ovchinnikova G, Ivanova NN, Kyrpides NC. 2016. Supporting community annotation and user collaboration in the integrated microbial genomes (IMG) system. BMC Genomics 17:307.

73. Markowitz VM, Chen I-MA, Palaniappan K, Chu K, Szeto E, Pillay M, Ratner A, Huang J, Woyke T, Huntemann M, Anderson I, Billis K, Varghese N, Mavromatis K, Pati A, Ivanova NN, Kyrpides NC. 2014. IMG 4 version of the integrated microbial genomes comparative analysis system. Nucleic Acids Research 42:D560–D567.

74. Katoh K, Standley DM. 2013. MAFFT Multiple Sequence Alignment Software Version 7: Improvements in Performance and Usability. Mol Biol Evol 30:772–780.

75. Katoh K, Misawa K, Kuma K, Miyata T. 2002. MAFFT: a novel method for rapid multiple sequence alignment based on fast Fourier transform. Nucleic Acids Res 30:3059–3066.

76. Minh BQ, Nguyen MAT, von Haeseler A. 2013. Ultrafast Approximation for Phylogenetic Bootstrap. Molecular Biology and Evolution 30:1188–1195.

77. Minh BQ, Schmidt HA, Chernomor O, Schrempf D, Woodhams MD, von Haeseler A, Lanfear R. 2020. IQ-TREE 2: New Models and Efficient Methods for Phylogenetic Inference in the Genomic Era. Molecular Biology and Evolution 37:1530–1534.

78. Mirdita M, Schütze K, Moriwaki Y, Heo L, Ovchinnikov S, Steinegger M. 2022. ColabFold: making protein folding accessible to all. Nat Methods 19:679–682.

79. Guzenko D, Burley SK, Duarte JM. 2020. Real time structural search of the Protein Data Bank. PLOS Computational Biology 16:e1007970.

80. Eisenberg D, Schwarz E, Komaromy M, Wall R. 1984. Analysis of membrane and surface protein sequences with the hydrophobic moment plot. J Mol Biol 179:125–142.

