## Supplemental Information for "A sterol reductase responsible for the unusual 8(14)-unsaturation in bacterial sterol production and degradation"

**This file includes:**

Figures S1-S9
Tables S1-S5
SI References

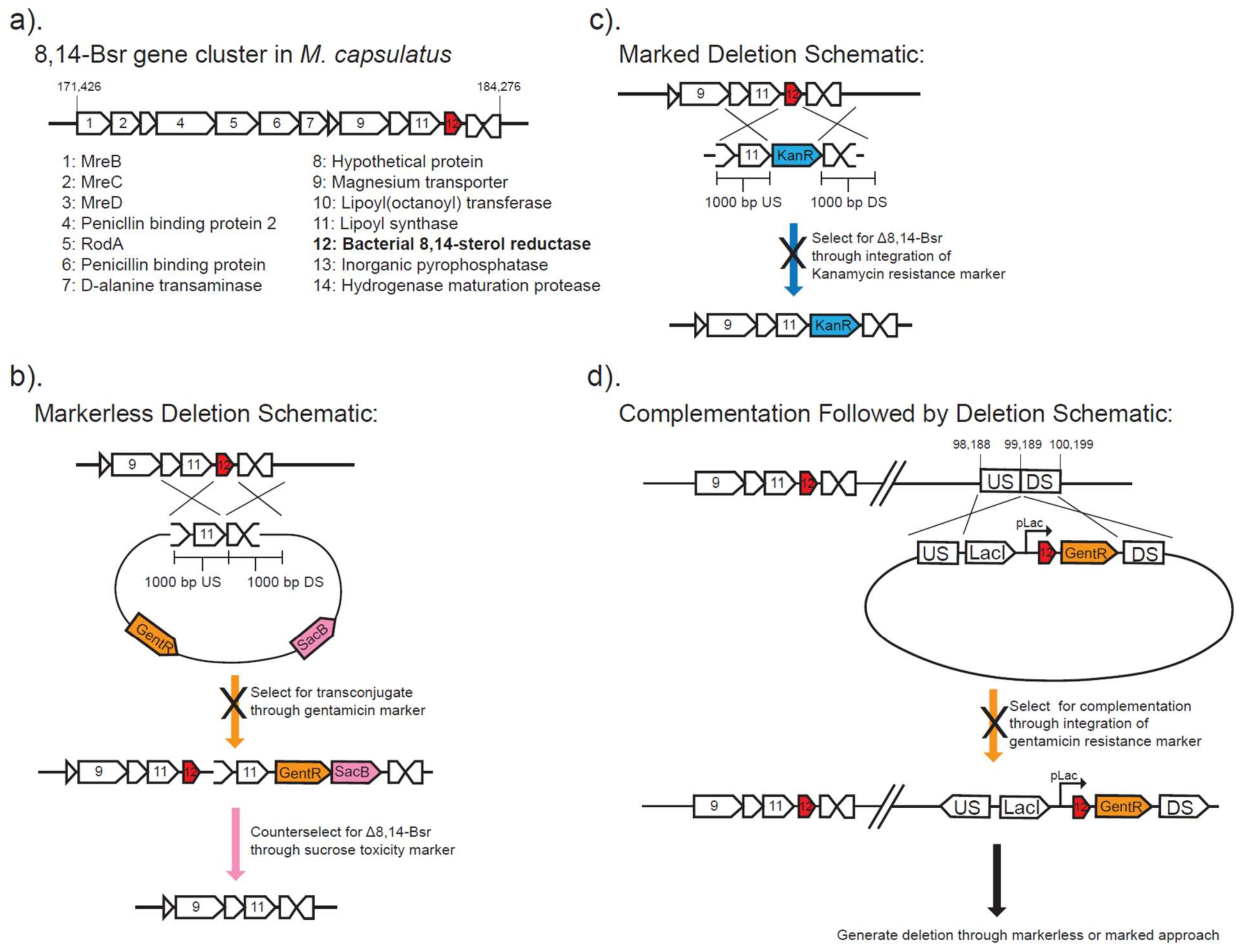


**Figure S1. Representative schematic of genetic approaches for attempted 8,14-Bsr gene deletion in *Methylococcus capsulatus* Texas.** a). The 8,14-Bsr homolog in *M. capsulatus* sits in a large gene cluster with genes likely involved in essential processes including cellular metabolism and cell shape. b). Our initial attempts at deleting 8,14-Bsr in *M. capsulatus* involved constructing a vector with flanking upstream and downstream regions and both a positive (gentamycin resistance) and negative selection (sucrose toxicity) marker. The vector was introduced through conjugation, and we attempted to select for integration of this vector into the chromosome through gentamycin resistance but did not recover gentamycin resistant clones. c). We next attempted to generate a marked deletion by flanking a kanamycin resistance marker with upstream and downstream regions of the 8,14-Bsr gene through overlap extension PCR. This PCR product was introduced to the bacterium through electroporation. We have had previous success deleting sterol biosynthesis genes in *M. capsulatus* Texas through this approach (1), however we were unable to recover kanamycin resistant clones. d). Finally, we tried to introduce 8,14-Bsr to a secondary chromosomal location under the expression of a lac promoter so we could then try to delete the original copy either through a marked or markerless approach, which might suggest essentiality of this gene product. Again, we have previously successfully integrated and expressed sterol biosynthesis genes from this location in *M. capsulatus* Texas (1), but here were unable to recover gentamycin resistant clones.

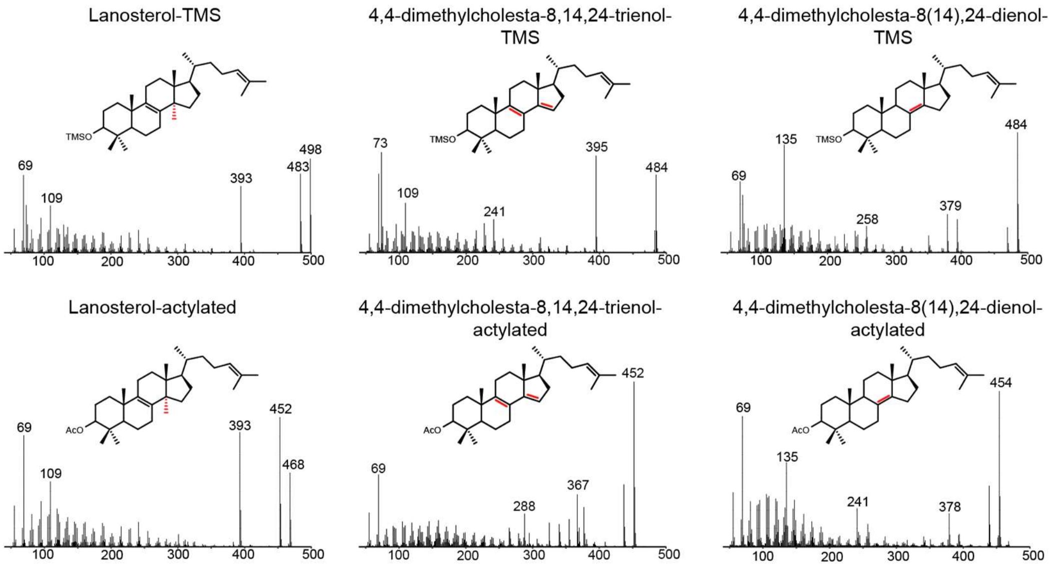


**Figure S2. Spectra of sterols identified in heterologous expression cultures.** Sterols were derivatized to trimethylsilyl (TMS) ether or acetyl groups and separated using an Agilent 7890B series gas chromatograph with a 60m Agilent DB17 column (60m x 0.25 mm i.d x 0.1 µm film thickness). Helium gas was used as a carrier and coupled to a 5977A series mass spectrometer. Sterols were identified based on elution time and spectra compared to published spectra in the NIST database or through NMR analysis. See Methods for full extraction and GC-MS details.

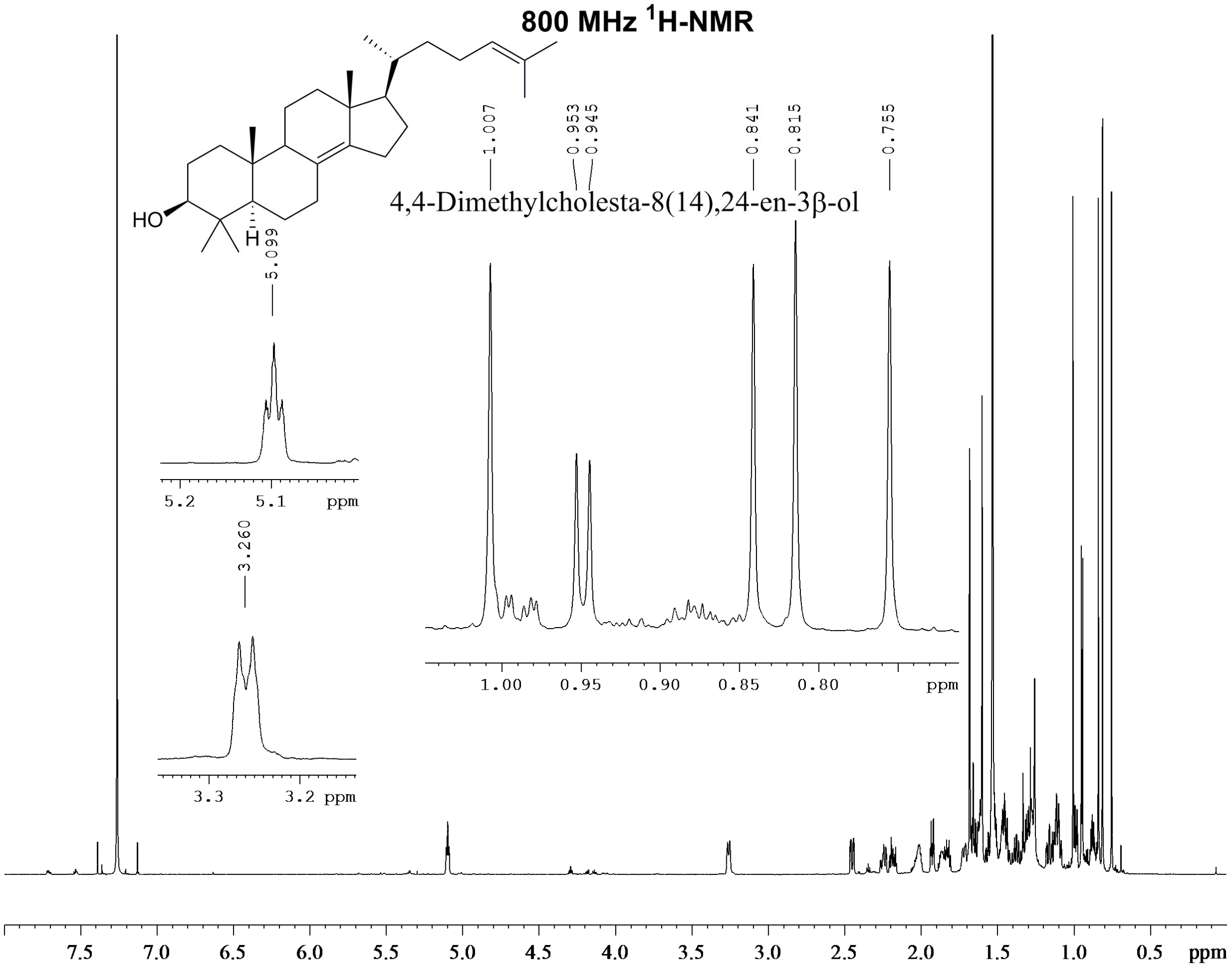


**Figure S3. $^1$H-NMR spectra of 4,4-dimethylcholesta-8(14), 24-dienol.**

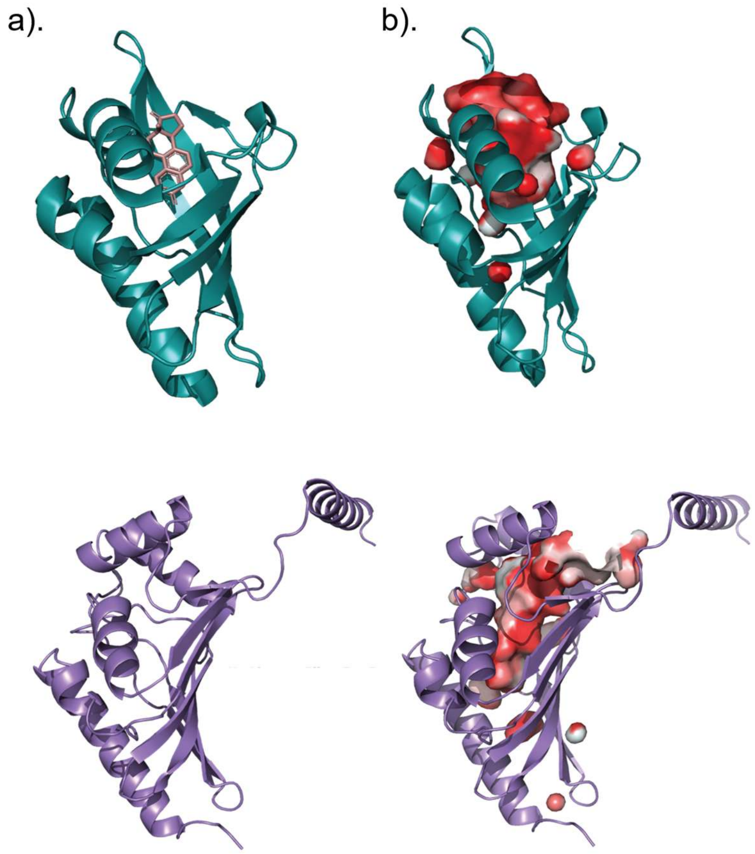


**Figure S4. Comparison of the modeled 8,14-Bsr homolog from *M. capsulatus* Texas to the crystalized structure of ketosteroid isomerase (PBD: 1OH0) from *Pseudomonas putida*.** a) Ribbon diagram of ketosteroid isomerase (teal) with inhibitor equilenin bound in the active site (2). The modeled 8,14-Bsr homolog (lilac) shares the same cone shaped fold as Ksi, but with additional alpha helices at the C-terminus. b) Hydrophobic active site pockets in ketosteroid isomerase (teal) and 8,14-Bsr (lilac). The hydrophobicity scale ranges from white for hydrophilic residues to red for hydrophobic residues and is based on the Eisenberg hydrophobicity scale (3). c) Characterized active site residues in ketosteroid isomerase (teal) and putative active site residues in 8,14-Bsr (lilac).

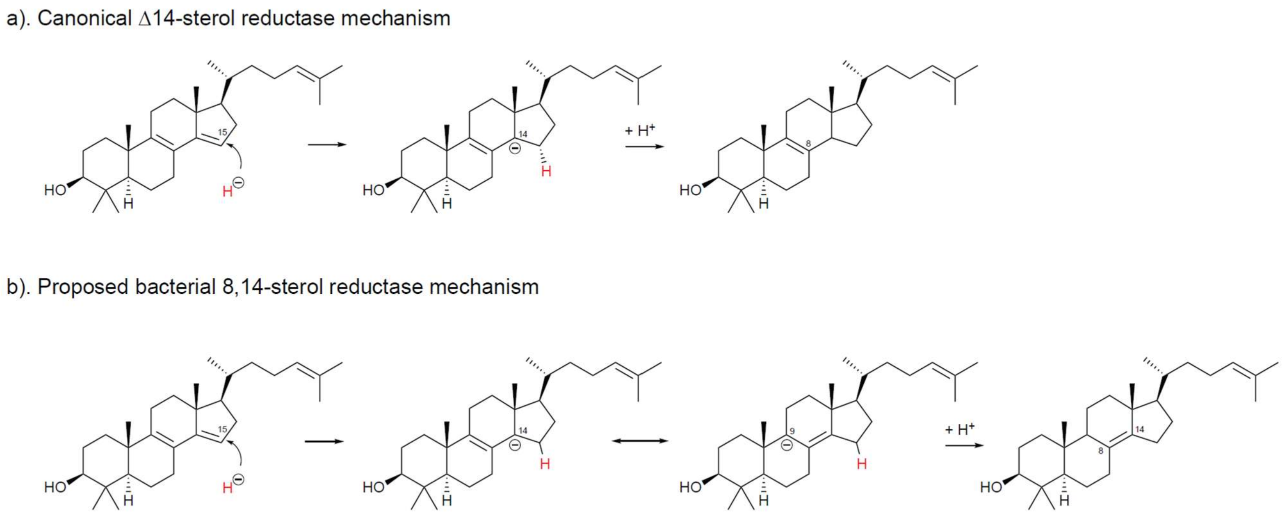


**Figure S5. A proposed mechanism for the bacterial 8,14-sterol reductase.** a). The canonical C-14 sterol reductase mechanism. A hydride from NADPH is added to the C-15 position. This generates an anion at C-14, which is then protonated to produce a Δ8 sterol. b). The proposed mechanism for 8,14-Bsr. Like the C-14 reductase, 8,14-Bsr adds hydride to its substrate at C-15, generating an anion at C-14. Through allylic rearrangement, this charge migrates to C-9, where it is deprotonated resulting in an 8(14) sterol.

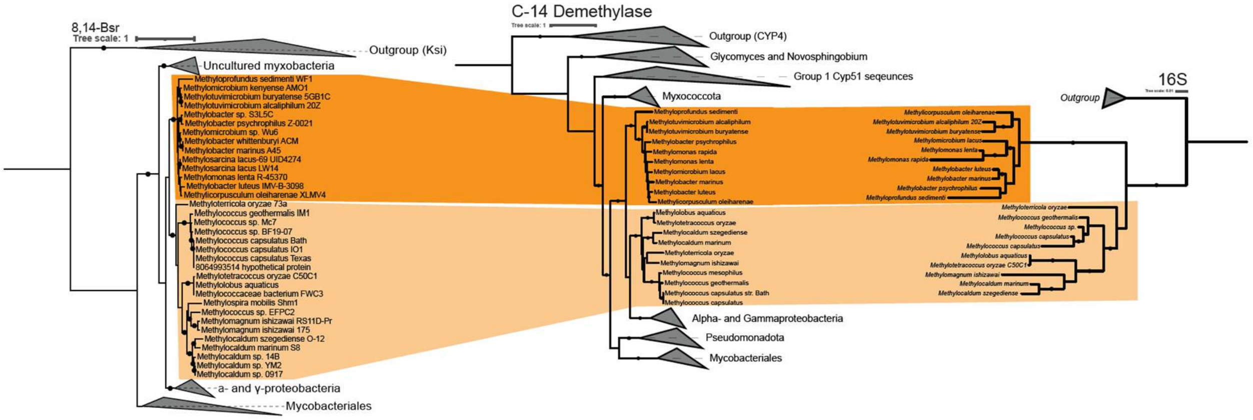


**Figure S6. Phylogenetic analysis of *Methylococcaceae* 8,14-Bsr and C-14 demethylase sequences in relation to taxonomic diversity.** A maximum likelihood trees of 16S sequences from *Methylococcaceae* species with *Pseudomonas* sequences as an outgroup, C-14 demethylase (Cyp51) sequences with Cyp4 sequences as an outgroup, and 8,14-Bsr sequences with ketosteroid isomerase (Ksi) sequences as an outgroup were generated using IQTree with the model of best fit and 5000 ultrafast bootstrap replicates. Branches with bootstrap support values >90 are denoted with a black circle. 8,14-Bsr, Cyp51 and 16S sequences largely cluster similarly.

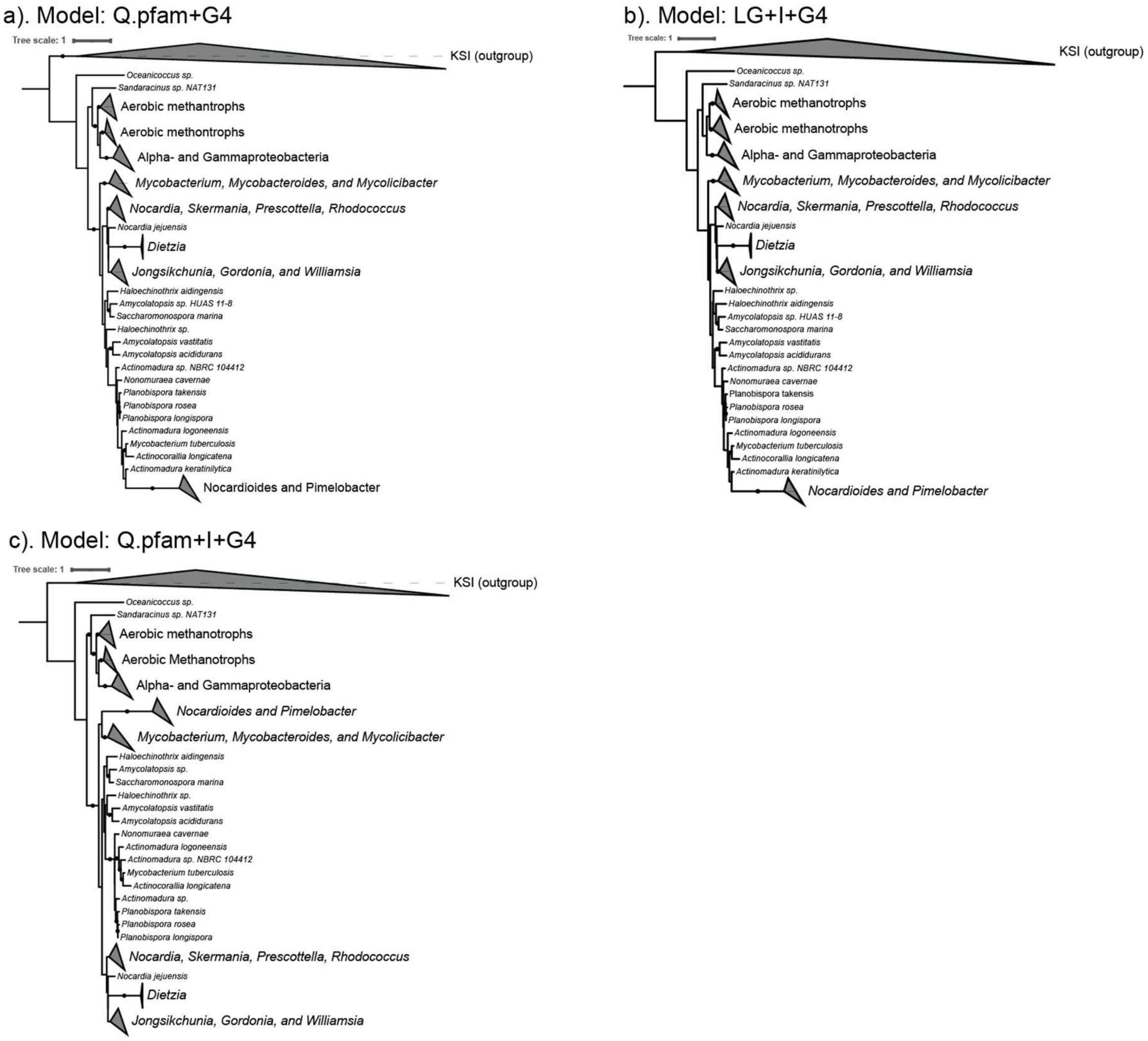


**Figure S7. Phylogenetic trees of the bacterial 8,14-sterol reductase (8,14-Bsr) generated using alternative models and parameters.** IQ-TREE was used to generate maximum likelihood trees with 5000 ultrafast bootstrap replicates and the following models: a) Q.pfam+G4, b) LG+I+G4, c) Q.pfam+I+G4. Branches with bootstrap support values greater than 90 are denoted with a black circle. Tree topology is largely maintained with some uncertainty in branching in the Mycobacteriales sequences.

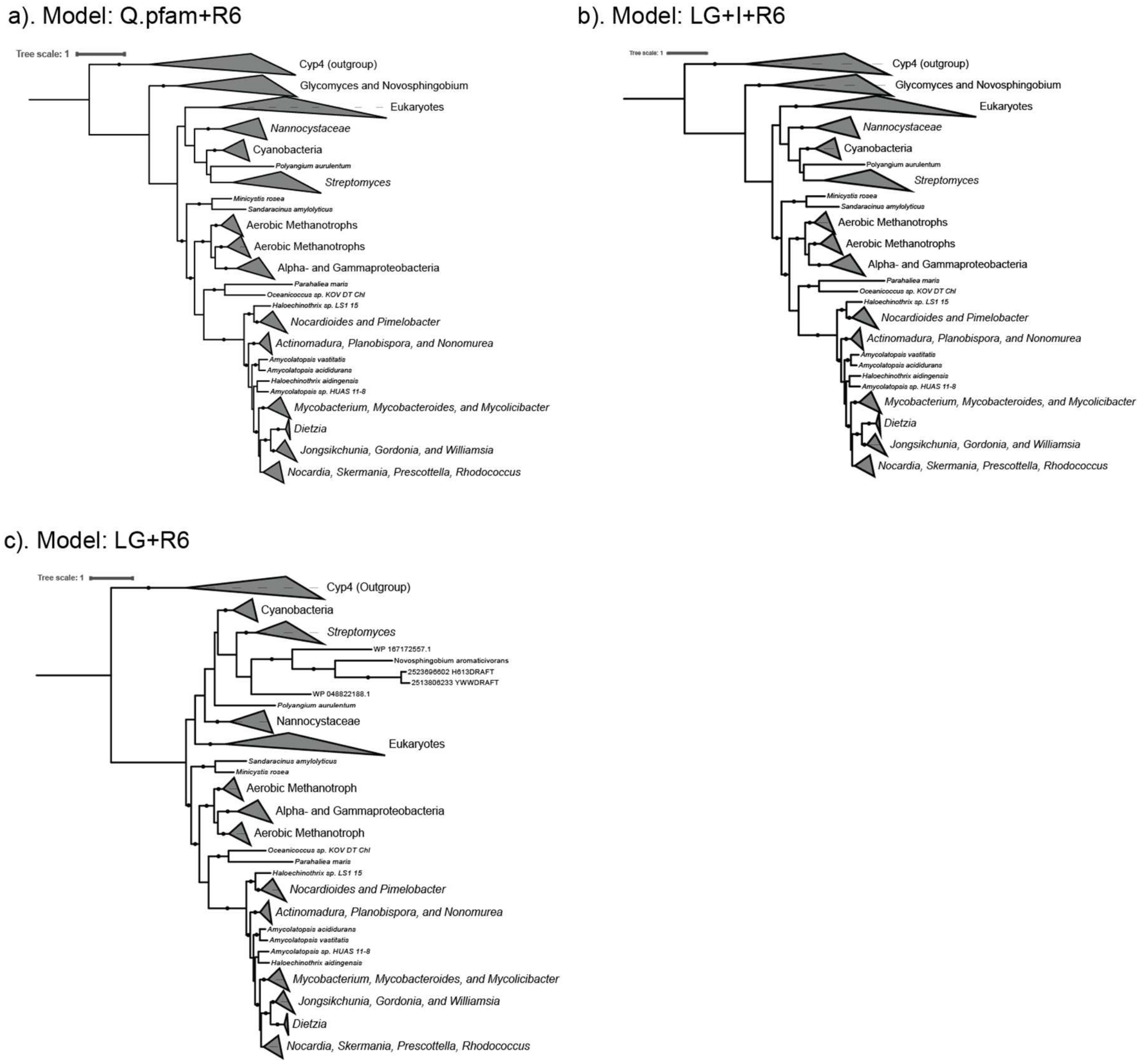


**Figure S8. Phylogenetic trees of the C-14 demethylase (CYP51) generated using alternative models and parameters.** IQ-TREE was used to generate maximum likelihood trees with 5000 ultrafast bootstrap replicates and the following models: a) Q.pfam+R6, b) LG+I+R6, c) LG+R6. Branches with bootstrap support values greater than 90 are denoted with a black circle. General tree topology is maintained regardless of the model used.

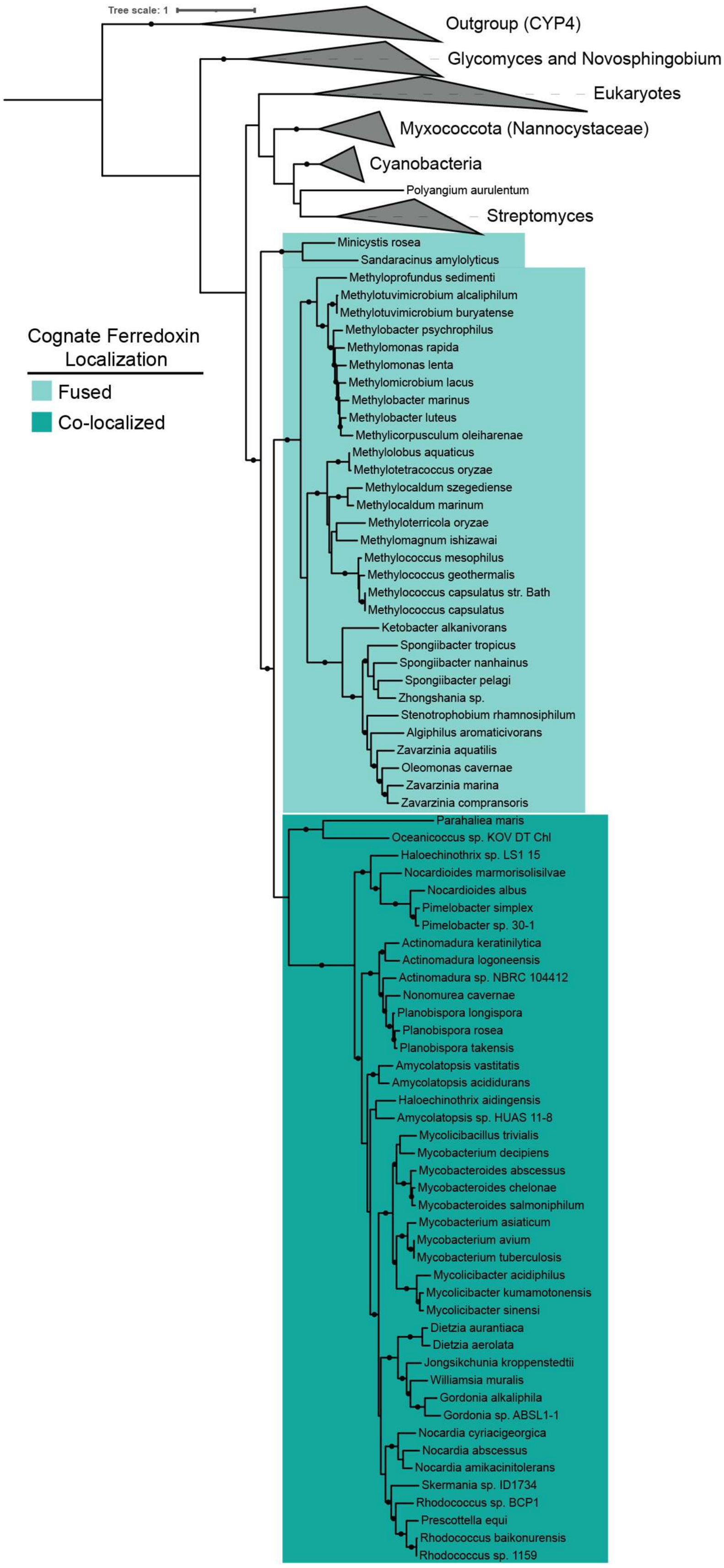


**Figure S9. C-14 demethylase sequences from bacteria that harbor bacterial 8,14-sterol reductase are fused or co-localized to their cognate ferredoxin.** A maximum likelihood trees of the C-14 demethylase (Cyp51) with Cyp4 as an outgroup were generated using IQTree with the model of best fit and 5000 ultrafast bootstrap replicates. Branches with bootstrap support values >90 are denoted with a black circle. Cyp51 sequences fused to their cognate ferredoxin are highlighted in light cyan. Cyp51sequneces col-localized with their cognate ferredoxin are highlighted in cyan. This physical relationship between Cyp51 and its ferredoxin is found only in the clade of bacteria that also have the bacterial 8,14-sterol reductase.

**Table S1. Comparative genomics to identify potential genes involved in 8(14)-unsaturation generation.** Genes in *Methylocococcus capsulatus* Texas that are also present in *Methylobacter luteus, Methylomonas lenta,* and *Methyloprofundus sedimenti* and absent in *Crenothrix polyspora*, *Methylohalobius crimeensis*, and *Methylomonas methanica*. This list was generated using the IMG phylogenetic profiler tool with a minimum e-value of e-05 and 30% identity. The gene characterized in this study is bolded.

| Locus Tag | Gene Product Name |
|---|---|
| H156DRAFT_0063 | Arm domain-containing DNA-binding protein |
| H156DRAFT_0889 | Delta24-sterol reductase |
| H156DRAFT_1017 | Outer membrane protein |
| H156DRAFT_1328 | (2Fe-2S) ferredoxin |
| H156DRAFT_1441 | Uncharacterized protein (TIGR00725 family)/uncharacterized protein (TIGR00730 family) |
| H156DRAFT_1746 | Sterol 14alpha-demethylase |
| H156DRAFT_1904 | Methionine sulfoxide reductase heme-binding subunit |
| **H156DRAFT_2205** | **hypothetical protein** |
| H156DRAFT_2631 | Paraquat-inducible protein A |
| H156DRAFT_2654 | Glycine cleavage system P protein (glycine dehydrogenase) subunit 2 |

**Table S2. Eukaryotes that produce de novo 8(14)-sterols have a C-14 reductase.** Using a BLASTp search of proteins or a tBLASTn search of transcriptomes (e-value e-05), we queried the genomes of eukaryotes reported to produce 8(14)-sterols for homologs to the canonical C-14 reductase. References to the literature reporting the sterol profile of these organisms are provided as well as the genbank accession number for the C-14 reductase homologs recovered by our BLAST searches.

| Organism | Sterol produced | C-14 reductase Accession Number | Reference |
|---|---|---|---|
| *Caenorhabditis elegans* | 4-methyl cholest-8(14)-enol | Not Applicable* | (4) |
| *Karenia brevis* | Gymnodinosterol | GFLM01054842 | (5) |
| *Amphidinium carterae* | Amphisterol | HBNO01050105 | (6) |
| *Testudodinium testudo* | Gymnodinosterol | No genome available | (7) |
| *Dvsidea herbacea* | 3-oxo-4,6,8(14)-steroids | No genome available | (8) |

*Does not synthesize sterols de novo but instead modifies exogenously acquired sterols that have already been demethylated at C-14.

**Table S3. Oligonucleotide primers used in this study.** MCAT indicates *Methylococcus capsulatus* Texas. Fwd indicates forward primer. Rv indicates reverse primer. Cyp51 indicates the C-14 demethylase. MCA0111 and H156DRAFT_2205 indicate the bacterial 8,14-sterol reductase. SDM indicates site directed mutagenesis primer.

| Oligonucleotide | Sequence | Notes |
|---|---|---|
| AL134 | TTGGGCTAGCAGGAGGAATTCAC ATGAGCCATCCCCCTTC | SLIC pBAD-NcoI-MCAT Cyp51 Fwd |
| AL135 | GACTCTAGAGGATCCCCGGGTACCTAATCCTC GCCCGAAGG | SLIC pBAD-NcoI-MCAT Cyp51 Rv |
| AL23 | CGTCACACTTTGCTATGCCA | pBAD sequencing primer Fwd |
| AL22 | CCGCCAGGCAAATTCTGTTT | pBAD sequencing primer Rv |
| AL373 | CCACCGCGGTGGCGGCCGCTATGCCCGCC TACCCC | SLIC pBAD-XbaI-MCAT MCA0111 Fwd |
| AL374 | GCCCGGGGGATCCACTAGTT TCAGCCGCCCGACAG | SLIC pBAD-XbaI-MCAT MCA0111 Rv |
| AL375 | TCGCTCGGTGCGATGGCTACCGAAGACGCC GA | SDM primer H156DRAFT_2205 Y37A |
| AL376 | ACCGAAGACGCCGAAGCTGGCTGGAACATG GGG | SDM primer H156DRAFT_2205 Y43A |
| AL377 | TTCGAGCAATGGCGCGCTCCGTACGACCACG TG | SDM primer H156DRAFT_2205 Y78A |
| AL378 | CAATGGCGCTACCCGGCTGACCACGTGGTGA TC | SDM primer H156DRAFT_2205 Y80A |
| AL379 | TGGAGCTGGCAGCGCGCTTTCTTCGACCTGG GC | SDM primer H156DRAFT_2205 D131A |
| AL380 | TCGCTCGGTGCGATGTTCACCGAAGACGCCG A | SDM primer H156DRAFT_2205 Y37F |
| AL381 | ACCGAAGACGCCGAATTCGGCTGGAACATGG GG | SDM primer H156DRAFT_2205 Y43F |
| AL382 | TTCGAGCAATGGCGCTTCCCGTACGACCACG TG | SDM primer H156DRAFT_2205 Y78F |
| AL383 | CAATGGCGCTACCCGTTCGACCACGTGGTGA TC | SDM primer H156DRAFT_2205 Y80F |
| AL384 | CAGCGCGATTTCTTCGCTCTGGGCAATGTCAA G | SDM primer H156DRAFT_2205 D133A |

**Table S4. Plasmids used in this study.** (*) indicates this plasmid was generated in this study, RBS indicates ribosome binding site, MCAT indicates *Methylococcus capsulatus* Texas. MCA0111 corresponds to the bacterial 8,14-sterol reductase. Cyp51 corresponds to the C-14 demethylase.

| Plasmid | Description | Reference |
|---|---|---|
| pTrc-sqs-synRBS-osc-synRBS-smo (pABB501) | MEALZ_3096-MEALZ_0768-MEALZ_0767 (Squalene synthase-Oxidosqualene cyclase-Squalene epoxidase) optimized expression plasmid (altered osc and smo RBSs). | (1) |
| pBAD1031K (pABB466) | pRV1031 ori, pBAD promoter, Kanr | (9) |
| pBAD1031K_MCAT_Cyp51 (pAL7018) | H156DRAFT_1746 (Cyp51) expression plasmid<br><br>H156DRAFT_2205 was amplified by PCR with primers AL134 and AL135. The fragment was assembled by SLIC into the NcoI site of pABB466. Sequence was confirmed with oligos AL22 and AL23. | (*) |
| pBAD1031K_MCAT_MCA0111 (pAL7406) | H156DRAFT_2205 (MCA0111) expression plasmid<br><br>H156DRAFT_2205 was amplified by PCR with primers AL373 and AL374. The fragment was assembled by SLIC into the XbaI site of pABB66. Sequence was confirmed with oligos AL22 and AL23. | (*) |
| pBAD1031K_MCAT_Cyp51_MCA0111 (pAL7019) | H156DRAFT_1746 (Cyp51) and 2205 (MCA0111) expression plasmid<br><br>H156DRAFT_2205 was amplified by PCR with primers AL373 and AL374. The fragment was assembled by SLIC into the XbaI site of pAL7018. Sequence was confirmed with oligos AL22 and AL23. | (*) |
| pBAD1031K_MCAT_Cyp51_MCA0111_Y37A (AL7418) | H156DRAFT_2205_Y37A mutagenesis expression plasmid<br><br>H156DRAFT was mutated through amplification of pAL7019 with primer AL375. Sequence was confirmed with oligos AL22 and AL23. | (*) |

| | | |
|---|---|---|
| pBAD1031K_MCAT_<br>Cyp51_MCA0111_Y43_A (AL7419) | H156DRAFT_2205_Y43A mutagenesis expression plasmid<br><br>H156DRAFT was mutated through amplification of pAL7019 with primer AL376. Sequence was confirmed with oligos AL22 and AL23. | (*) |
| pBAD1031K_MCAT_<br>Cyp51_MCA0111_Y78A (AL7420) | H156DRAFT_2205_Y78A mutagenesis expression plasmid<br><br>H156DRAFT was mutated through amplification of pAL7019 with primer AL377. Sequence was confirmed with oligos AL22 and AL23. | (*) |
| pBAD1031K_MCAT_<br>Cyp51_MCA0111_Y80A (AL7421) | H156DRAFT_2205_Y80A mutagenesis expression plasmid<br><br>H156DRAFT was mutated through amplification of pAL7019 with primer AL378. Sequence was confirmed with oligos AL22 and AL23. | (*) |
| pBAD1031K_MCAT_<br>Cyp51_MCA0111_D131A (AL7422) | H156DRAFT_2205_D131A mutagenesis expression plasmid<br><br>H156DRAFT was mutated through amplification of pAL7019 with primer AL379. Sequence was confirmed with oligos AL22 and AL23. | (*) |
| pBAD1031K_MCAT_<br>Cyp51_MCA0111_D133A (AL7444) | H156DRAFT_2205_D131A mutagenesis expression plasmid<br><br>H156DRAFT was mutated through amplification of pAL7019 with primer AL383. Sequence was confirmed with oligos AL22 and AL23. | (*) |
| pBAD1031K_MCAT_<br>Cyp51_MCA0111_Y37F (AL7423) | H156DRAFT_2205_Y37F mutagenesis expression plasmid<br><br>H156DRAFT was mutated through amplification of pAL7019 with primer AL380. Sequence was confirmed with oligos AL22 and | (*) |

| | | |
|---|---|---|
| | AL23. | |
| pBAD1031K_MCAT_ Cyp51_MCA0111_Y43F (AL7424) | H156DRAFT_2205_Y43F mutagenesis expression plasmid<br><br>H156DRAFT was mutated through amplification of pAL7019 with primer AL381. Sequence was confirmed with oligos AL22 and AL23. | (*) |
| pBAD1031K_MCAT_ Cyp51_MCA0111_Y78F (pAL7425) | H156DRAFT_2205_Y78F mutagenesis expression plasmid<br><br>H156DRAFT was mutated through amplification of pAL7019 with primer AL381. Sequence was confirmed with oligos AL22 and AL23. | (*) |
| pBAD1031K_MCAT_ Cyp51_MCA0111_Y80F (pAL7426) | H156DRAFT_2205_Y80F mutagenesis expression plasmid<br><br>H156DRAFT was mutated through amplification of pAL7019 with primer AL382. Sequence was confirmed with oligos AL22 and AL23. | (*) |
| pBAD1031K_MCAT_MCA0111_Y37A (AL7427) | H156DRAFT_2205_Y37A mutagenesis expression plasmid<br><br>H156DRAFT was mutated through amplification of pAL7406 with primer AL375. Sequence was confirmed with oligos AL22 and AL23. | (*) |
| pBAD1031K_MCAT_ MCA0111_Y43_A (AL7428) | H156DRAFT_2205_Y43A mutagenesis expression plasmid<br><br>H156DRAFT was mutated through amplification of pAL7406 with primer AL376. Sequence was confirmed with oligos AL22 and AL23. | (*) |
| pBAD1031K_MCAT_ MCA0111_Y78A (AL7429) | H156DRAFT_2205_Y78A mutagenesis expression plasmid<br><br>H156DRAFT was mutated through amplification of pAL7406 with primer AL377. Sequence was | (*) |

| | | |
|---|---|---|
| | confirmed with oligos AL22 and AL23. | |
| pBAD1031K_MCAT_ MCA0111_Y80A (AL7430) | H156DRAFT_2205_Y80A mutagenesis expression plasmid<br><br>H156DRAFT was mutated through amplification of pAL7406 with primer AL378. Sequence was confirmed with oligos AL22 and AL23. | (*) |
| pBAD1031K_MCAT_ MCA0111_D131A (AL7431) | H156DRAFT_2205_D131A mutagenesis expression plasmid<br><br>H156DRAFT was mutated through amplification of pAL7406 with primer AL379. Sequence was confirmed with oligos AL22 and AL23. | (*) |
| pBAD1031K_MCAT_ MCA0111_D133A (AL7445) | H156DRAFT_2205_D131A mutagenesis expression plasmid<br><br>H156DRAFT was mutated through amplification of pAL7406 with primer AL383. Sequence was confirmed with oligos AL22 and AL23. | (*) |
| pBAD1031K_MCAT_ MCA0111_Y37F (AL7432) | H156DRAFT_2205_Y37F mutagenesis expression plasmid<br><br>H156DRAFT was mutated through amplification of pAL7406 with primer AL380. Sequence was confirmed with oligos AL22 and AL23. | (*) |
| pBAD1031K_MCAT_ MCA0111_Y43F (AL7433) | H156DRAFT_2205_Y43F mutagenesis expression plasmid<br><br>H156DRAFT was mutated through amplification of pAL7406 with primer AL381. Sequence was confirmed with oligos AL22 and AL23. | (*) |
| pBAD1031K_MCAT_ MCA0111_Y78F (pAL7434) | H156DRAFT_2205_Y78F mutagenesis expression plasmid<br><br>H156DRAFT was mutated through amplification of pAL7406 with | (*) |

| | | |
|---|---|---|
| | primer AL381. Sequence was confirmed with oligos AL22 and AL23. | |
| pBAD1031K_MCAT_ MCA0111_Y80F (pAL7435) | H156DRAFT_2205_Y80F mutagenesis expression plasmid<br><br>H156DRAFT was mutated through amplification of pAL7406 with primer AL382. Sequence was confirmed with oligos AL22 and AL23. | (*) |

**Table S5. Heterologous expression strains used in this study.** All expression strains are *E. coli* DH10B with pJBEI2997 (pABB302, CmR), pTrc (pABB278 or derivatives, AmpR), pSRK (pABB492 or derivatives, Gmr) (where indicated), and pBAD1031K (pABB466 or derivatives, KanR) (where indicated).

| Expression Strain | Plasmids |
|---|---|
| PVW 7011 | pJBEI2997 (pABB302), pTrc-sqs-synRBS-osc-synRBS-smo (pABB501), pSRKGm-lacUV5-rbs5 (pABB492), pBAD1031K (pABB466) |
| PVW 7020 | pJBEI2997 (pABB302), pTrc-sqs-synRBS-osc-synRBS-smo (pABB501), pBAD1031K-Cyp51 (pAL7018) |
| PVW 7022 | pJBEI2997 (pABB302), pTrc-sqs-synRBS-osc-synRBS-smo (pABB501), pBAD1031K-Cyp51-MCA0111 (pAL7019) |
| PVW 7408 | pJBEI2997 (pABB302), pTrc-sqs-synRBS-osc-synRBS-smo (pABB501), pSRKGm-lacUV5-rbs5-Cyp51 (pAL7158), pBAD1031K-MCA0111 (pAL7406) |
| PVW 7409 | pJBEI2997 (pABB302), pTrc-sqs-synRBS-osc-synRBS-smo (pABB501), pBAD1031K-Cyp51-MCA0111-Y37A (pAL7418) |
| PVW 7410 | pJBEI2997 (pABB302), pTrc-sqs-synRBS-osc-synRBS-smo (pABB501), pBAD1031K-Cyp51-MCA0111-Y43A (pAL7419) |
| PVW 7411 | pJBEI2997 (pABB302), pTrc-sqs-synRBS-osc-synRBS-smo (pABB501), pBAD1031K-Cyp51-MCA0111-Y78A (pAL7420) |
| PVW 7412 | pJBEI2997 (pABB302), pTrc-sqs-synRBS-osc-synRBS-smo (pABB501), pBAD1031K-Cyp51-MCA0111-Y80A (pAL7421) |
| PVW 7413 | pJBEI2997 (pABB302), pTrc-sqs-synRBS-osc-synRBS-smo (pABB501), pBAD1031K-Cyp51-MCA0111-D131A (pAL7422) |
| PVW 7414 | pJBEI2997 (pABB302), pTrc-sqs-synRBS-osc-synRBS-smo (pABB501), pBAD1031K- Cyp51-MCA0111-Y37F (pAL7423) |
| PVW 7415 | pJBEI2997 (pABB302), pTrc-sqs-synRBS-osc-synRBS-smo (pABB501), pBAD1031K- Cyp51-MCA0111-Y43F (pAL7424) |
| PVW 7416 | pJBEI2997 (pABB302), pTrc-sqs-synRBS-osc-synRBS-smo (pABB501), pBAD1031K- Cyp51-MCA0111-Y78F (pAL7425) |
| PVW 7417 | pJBEI2997 (pABB302), pTrc-sqs-synRBS-osc-synRBS-smo (pABB501), pBAD1031K- Cyp51-MCA0111-Y80F (pAL7426) |
| PVW 7436 | pJBEI2997 (pABB302), pTrc-sqs-synRBS-osc-synRBS-smo (pABB501), pBAD1031K-MCA0111-Y37A (pAL7427) |
| PVW 7437 | pJBEI2997 (pABB302), pTrc-sqs-synRBS-osc-synRBS-smo (pABB501), pBAD1031K-MCA0111-Y43A (pAL7428) |
| PVW 7438 | pJBEI2997 (pABB302), pTrc-sqs-synRBS-osc-synRBS-smo (pABB501), pBAD1031K-MCA0111-Y78A (pAL7429) |
| PVW 7439 | pJBEI2997 (pABB302), pTrc-sqs-synRBS-osc-synRBS-smo (pABB501), pBAD1031K-MCA0111-Y80A (pAL7430) |
| PVW 7423 | pJBEI2997 (pABB302), pTrc-sqs-synRBS-osc-synRBS-smo (pABB501), pBAD1031K-MCA0111-D131A (pAL7431) |
| PVW 7440 | pJBEI2997 (pABB302), pTrc-sqs-synRBS-osc-synRBS-smo (pABB501), pBAD1031K- MCA0111-Y37F (pAL7432) |
| PVW 7441 | pJBEI2997 (pABB302), pTrc-sqs-synRBS-osc-synRBS-smo (pABB501), pBAD1031K-MCA0111-Y43F (pAL7433) |
| PVW 7442 | pJBEI2997 (pABB302), pTrc-sqs-synRBS-osc-synRBS-smo (pABB501), pBAD1031K-MCA0111-Y78F (pAL7434) |

| | |
|---|---|
| PVW 7443 | pJBEI2997 (pABB302), pTrc-sqs-synRBS-osc-synRBS-smo (pABB501), pBAD1031K- MCA0111-Y80F (pAL7435) |
| PVW 7446 | pJBEI2997 (pABB302), pTrc-sqs-synRBS-osc-synRBS-smo (pABB501), pBAD1031K- Cyp51-MCA0111-D133A (pAL7444) |
| PVW 7447 | pJBEI2997 (pABB302), pTrc-sqs-synRBS-osc-synRBS-smo (pABB501), pBAD1031K- MCA0111-D133A (pAL7445) |

**References:**


1. Lee AK, Banta AB, Wei JH, Kiemle DJ, Feng J, Giner J-L, Welander PV. 2018. C-4 sterol demethylation enzymes distinguish bacterial and eukaryotic sterol synthesis. Proceedings of the National Academy of Sciences 115:5884–5889.

2. Kim SW, Cha S-S, Cho H-S, Kim J-S, Ha N-C, Cho M-J, Joo S, Kim KK, Choi KY, Oh B-H. 1997. High-Resolution Crystal Structures of Δ5-3-Ketosteroid Isomerase with and without a Reaction Intermediate Analogue. Biochemistry 36:14030–14036.

3. Eisenberg D, Schwarz E, Komaromy M, Wall R. 1984. Analysis of membrane and surface protein sequences with the hydrophobic moment plot. J Mol Biol 179:125–142.

4. Chitwood DJ, Lusby WR, Lozano R, Thompson MJ, Svoboda JA. 1984. Sterol metabolism in the nematodeCaenorhabditis elegans. Lipids 19:500–506.

5. Giner J-L, Faraldos JA, Boyer GL. 2003. Novel sterols of the toxic dinoflagellate karenia brevis (dinophyceae): a defensive function for unusual marine sterols?1. Journal of Phycology 39:315–319.

6. Withers NW, Goad LJ, Goodwin TW. 1979. A new sterol, 4α-methyl-5α-ergosta-8(14),24(28)-dien-3β-ol, from the marine dinoflagellate *Amphidinium carterae*. Phytochemistry 18:899–901.

7. Leblond JD, Elkins LC, Sabir K, Graeff JE. 2023. Sterols of Testudodinium testudo (formerly Amphidinium testudo): Production of the Δ8(14) sterol gymnodinosterol and chemotaxonomic relationship to the Kareniaceae. Journal of Eukaryotic Microbiology 70:e12929.

8. Kobayashi M, Krishna MM, Ishida K, Anjaneyulu V. 1992. Marine Sterols. XXII. Occurrence of 3-Oxo-4, 6, 8(14)-triunsaturated Steroids in the Sponge Dvsidea herbacea. Chem Pharm Bull 40:72–74.

9. Chakravartty V, Cronan JE. 2015. A series of medium and high copy number arabinose-inducible *Escherichia coli* expression vectors compatible with pBR322 and pACYC184. Plasmid 81:21–26.
